# A near-infrared fluorescent genetically encoded calcium ion biosensor with a direct response

**DOI:** 10.64898/2026.11.15.766740

**Authors:** Marina Musa, Rina Hashizume, Takumi Kobayashi, Kenwa Tamagawa, Rintaro Udo, Shosei Imai, Kazuhiro Sawada, Osamu Nureki, Yusuke Nasu, Takuya Terai, Robert E. Campbell, Misao Mizuno, Yong Qian

**Author notes:** These authors contributed equally to this work (Marina Musa, Rina Hashizume, Takumi Kobayashi). Corresponding authors. (T.T.); (R.E.C.); (M. Mizuno, for correspondence related to Raman spectroscopy); (Y.Q.).

## Abstract

Near-infrared genetically encoded calcium ion (Ca^2+^) indicators (NIR GECIs) enable Ca^2+^ imaging at longer wavelengths than green or red fluorescent GECIs, providing reduced autofluorescence and phototoxicity, and improved compatibility with deep-tissue and multiplexed imaging applications. We have previously reported biliverdin (BV)-binding phytochrome-based NIR GECIs such as NIR-GECO2G that exhibit inverse fluorescence responses. Although inverse response GECIs can effectively report Ca^2+^ dynamics, their high baseline fluorescence in the resting apo state causes faster photobleaching and substantial out-of-focus background fluorescence, complicating signal interpretation. Here, using the inverse response NIR-GECO2G as the starting scaffold, we developed a direct response Ca^2+^ indicator, designated dNIR-GECO1.0, through iterative rounds of protein engineering and screening. dNIR-GECO1.0 exhibited greater molecular brightness than NIR-GECO2G, robust fluorescence in mammalian cells without exogenous BV supplementation, and a high dynamic range for reporting Ca^2+^ changes. A detailed mechanistic investigation identified the key mutations within the Ca^2+^ binding domain that were necessary to convert the inverse response indicator into a direct response indicator. Resonance Raman spectroscopic analysis provided insight into the specific changes in the BV chromophore resulting from the Ca^2+^-dependent conformational change.

## INTRODUCTION

Calcium ions (Ca^2+^) are involved in a wide range of physiological processes, including muscle contraction, enzyme activation, signal transduction, and neuronal activity^1,2^. Genetically encoded Ca^2+^ indicators (GECIs) enable optical visualization of Ca^2+^ dynamics with high spatiotemporal resolution and cell-type specificity, making them powerful tools for studying Ca^2+^ signaling in cultured cells and in model organisms. Over the past two decades, extensive engineering of green and red fluorescent protein-based sensors has produced highly optimized visible-spectrum (400-600 nm) GECIs with excellent sensitivity, kinetics, and brightness^3–8^. Despite their high performance, those GECIs still face challenges when used for deep-tissue imaging and multiplexed imaging with either optogenetic actuators or other fluorescent probes, where light scattering, tissue absorption, autofluorescence, and spectral crosstalk obscure their fluorescence signals^5^.

To address these limitations, there have been increasing efforts to develop far-red (600–650 nm) and near-infrared (NIR; 650–900 nm) fluorescent GECIs (NIR GECIs). Longer excitation and emission wavelengths would enable deeper tissue imaging by reducing light scattering, tissue absorption, and autofluorescence^9–13^. In addition, the spectral separation from visible fluorophores would facilitate multiplexed imaging with optogenetic actuators or other fluorescent reporters, allowing simultaneous control and visualization of multiple biological processes.

We have previously reported the NIR-GECO series of NIR GECIs based on NIR fluorescent proteins (NIR-FPs) derived from biliverdin (BV)-binding bacterial phytochromes. This series includes NIR-GECO1^14^, NIR-GECO2^15^, NIR-GECO2G^15^, and iBB-GECO^16^ based on mIFP^17^, and NIR-GECO3^18^ based on miRFP680^19^. Notably, all of these variants are inverse response indicators with fluorescence decreasing upon Ca^2+^ binding. For imaging of transient increases in Ca^2+^ concentration, as occur during cell signaling and neuronal activity, the cell spends most of its time in the low Ca^2+^ concentration state, which, in the case of inverse response GECIs, is the bright state. This bright state is generally more susceptible to photobleaching and the resulting baseline drift during long-term imaging. Additionally, this high baseline brightness causes cells situated outside the focal plane to emit continuous fluorescence, increasing background fluorescence in complex tissues.

Recently, some direct response NIR Ca^2+^ indicators such as GAF-CaMP2–sfGFP^11^ and GAF-CaMP3–sfGFP^12^ have been reported, in which fluorescence increases upon Ca^2+^ binding. However, these sensors are dim when expressed in mammalian cells and require supplementation with exogenous BV for reliable detection of fluorescence signals. In addition, their small responses to Ca^2+^ (maximum Δ*F*/*F* = 2.3 as purified protein) further limit their utility for cell imaging. As an alternative approach to BV-binding NIR-FPs, the NIR GECIs HaloCaMP and WHaloCaMP have been developed by inserting the Ca^2+^-sensing domain and peptide into the self-labeling protein HaloTag^20,21^. These indicators are relatively bright and photostable, and exhibit increased fluorescence upon Ca^2+^ binding. However, they are not fully genetically encoded and require the addition of rhodamine dye derivatives before imaging.

Here, we report the dNIR-GECO series of direct response, fully genetically encoded NIR Ca^2+^ indicators derived from the NIR-GECO2G scaffold (**Figure 1A(i)**). Through extensive protein engineering and multiple rounds of directed evolution, we converted the inverse response behavior of NIR-GECO2G into a direct response while improving brightness, dynamic range, and performance in mammalian cells. In addition, we systematically characterized the mutations responsible for this functional switch and investigated the molecular mechanisms underlying the conversion from an inverse to a direct response. This work expands the landscape of phytochrome-based biosensors and lays a foundation for developing further improved high-performance biosensors for Ca^2+^ and other analytes.

**Figure 1.**
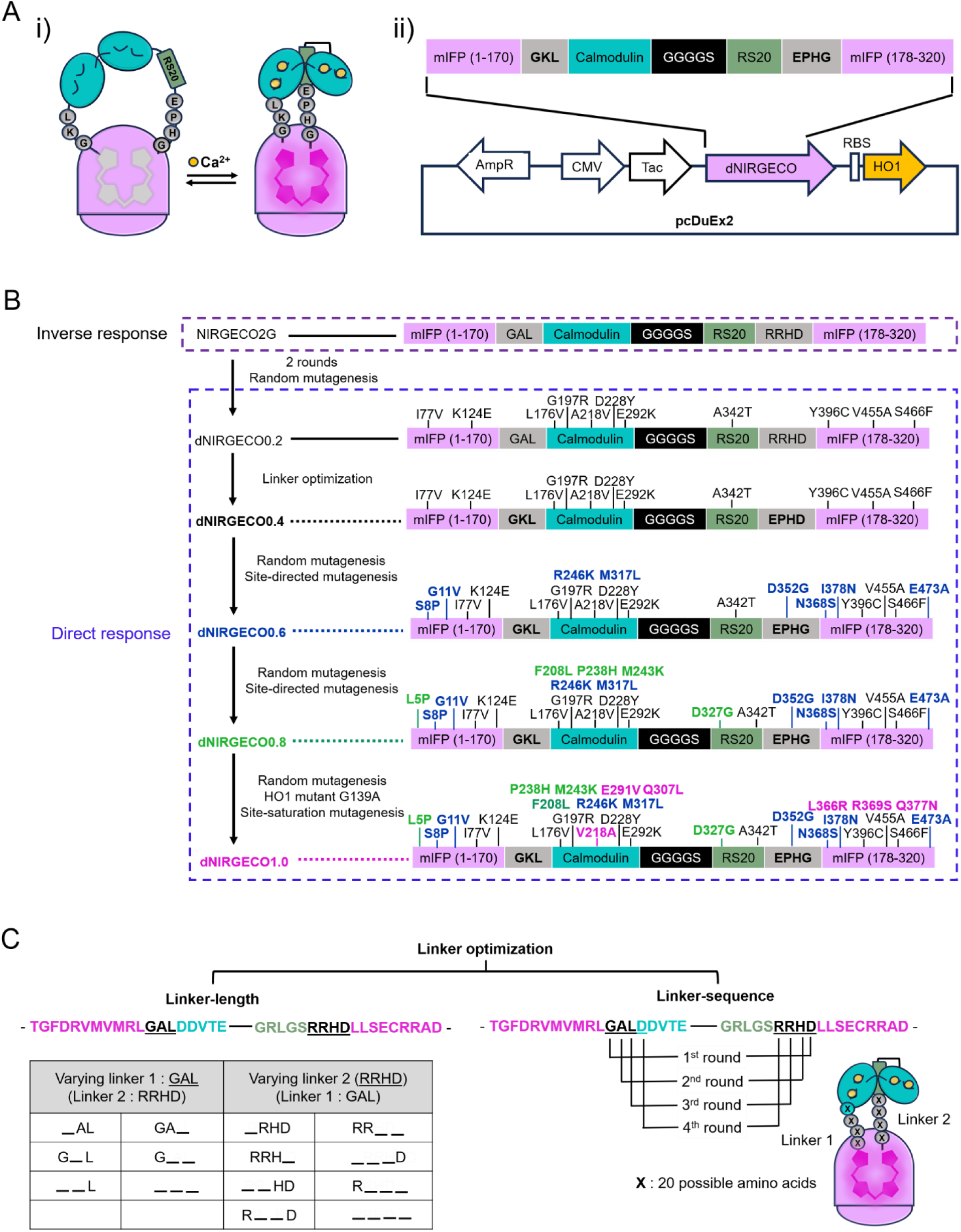
Design and development of dNIR-GECO1.0. (A) (i) Schematic representation of dNIR-GECO1.0. Calmodulin (CaM) is colored cyan, the RS20 peptide is colored sage green, and the m**I**FP scaffold is colored light magenta. The biliverdin (BV) chromophore in the middle is colored grey in the Ca^2+^-free state and dark magenta in the Ca^2+^-bound state, with calcium ions shown in yellow. (ii) Linear schematic presentation of dNIR-GECO1.0 expressed in the pcDuEx2 vector. (B) Overview of the directed evolution process leading to dNIR-GECO1.0. (C) Overview of the linker optimization strategies, including linker-length optimization by amino acid deletion and linker-sequence randomization.

## RESULTS

### Development of a direct response NIR GECI

In an effort to engineer an initial prototype NIR GECI with a direct response, we performed random mutagenesis on the whole gene of NIR-GECO2G and then cloned the mutated genes into our previously reported dual-expression vector pcDuEx2^14^ for expression in bacteria (**Figure 1A(ii)**). Bacteria lysates with expressed NIR-GECO variants were subjected to screening for positive fluorescence changes upon binding to Ca^2+^. After two rounds of screening, we identified a NIR-GECO variant exhibiting a direct response to Ca^2+^, which we named dNIR-GECO0.2 (**Figure 1B**). Compared with the parental NIR-GECO2G (Δ*F*/*F* = -6.6), dNIR-GECO0.2 showed a direct response with a Δ*F*/*F* of 1.05 but a much dimmer fluorescence (**Supplementary Table 1**). Sequence analysis revealed 11 amino acid substitutions compared with NIR-GECO2G, including six mutations within the CaM/RS20 sensing domain (Leu176Val, Gly197Arg, Ala218Val, Asp228Tyr, Glu292Lys and Ala342Thr) and five within mIFP domain (Ile77Val, Lys124Glu, Tyr396Cys, Val455Ala and Ser466Phe) (**Figure 1B, Supplementary Figure 1**).

We next tried to improve dNIR-GECO0.2 through linker optimization. The two linker regions were optimized independently by systematically deleting amino acid residues. Six variants of the first linker and eight variants of the second linker were generated, while keeping the other linker unchanged in each experiment (**Figure 1C**). However, none of these variants showed improved performance over dNIR-GECO0.2 (**Supplementary Figure 2**), suggesting that the original linker lengths were already optimal. We then performed linker sequence optimization using saturation mutagenesis, in which the amino acid sequences of the two linker regions, alongside with the first amino acid of the CaM domain, were randomized, and the generated variants were then screened for brightness and Ca^2+^ responses (**Figure 1C, Supplementary Table 2**). After four rounds of optimization, we identified dNIR-GECO0.4, which exhibited increased fluorescence intensity compared with dNIR-GECO0.2. To further improve dNIR-GECO0.4, eight additional rounds of directed evolution via random mutagenesis of the whole gene, followed by fluorescence-based screening, were performed. A number of beneficial mutations were identified, including Ser8Pro, Met317Leu, Asn368Ser, Ile378Asn, Glu473Ala, Gly11Val, Arg246Lys, and Asp352Gly. The resulting variant, dNIR-GECO0.6, exhibited a Δ*F*/*F* of approximately 8.0 in purified protein (**Supplementary Table 1**). We next expressed dNIR-GECO0.6 in HeLa cells to evaluate its performance in mammalian cells. Unfortunately, the fluorescence of dNIR-GECO0.6 was barely detectable when no exogenous BV was added, suggesting inefficient binding of dNIR-GECO0.6 to endogenous BV.

Given that external BV supplementation is undesirable for imaging applications, we further engineered dNIR-GECO variants for improved endogenous BV binding and cellular brightness. We performed additional rounds of directed evolution, including screening in purified proteins and mammalian cells. After seven rounds of evolution, we identified dNIR-GECO0.8, which contains five additional mutations (Leu5Pro, Phe208Leu, Pro238His, Met243Lys, and Asp327Gly). dNIR-GECO0.8 exhibited improved fluorescence response in purified protein (Δ*F*/*F* = 12.1) and, importantly, detectable fluorescence in HeLa cells without exogenous BV supplementation, although its cellular brightness remained relatively weak.

To further improve its cellular performance, we continued the evolution of dNIR-GECO0.8 in a total of nine rounds of random mutagenesis and site-saturation mutagenesis. The first three rounds yield several variants with exceptionally high Ca^2+^ responses in purified proteins, including High-Response Variant 1 (HRV1, Δ*F*/*F* = 38.2), HRV2 (Δ*F*/*F* = 58.8), and HRV3 (Δ*F*/*F* = 44.3). Their large dynamic ranges suggest that these variants could provide useful starting scaffolds for engineering NIR fluorescent indicators for other analytes. However, all three variants exhibited very weak baseline fluorescence in mammalian cells, limiting their practical use for Ca^2+^ imaging (**Supplementary Table 3)**. Therefore, we took a step back and used dNIR-GECO0.8 as the template again for evolution and performed another six rounds of screening with a focus on variants’ brightness in HeLa cells. These rounds of directed evolution included random mutagenesis and site-saturation mutagenesis targeting residues near the BV-binding pocket (**Supplementary Figure 3**). We also introduced the Ala218Val reversion, the original residue in NIR-GECO2G, which improved cellular brightness. To facilitate the screening for effective BV-binding, we used the Heme Oxygenase-1 (HO1) mutant Gly139Ala reported in our previous study^18^ during the final stage of directed evolution. Ultimately, these efforts resulted in dNIR-GECO1.0, which showed improved performance in both purified protein and in HeLa cells (**Supplementary Table 1**), with a total of 28 amino acid substitutions compared to the parental NIR-GECO2G sequence (**Supplementary Figure 1**).

### *In vitr*o characterization of dNIR-GECO1.0 as purified protein

We next characterized the evolved dNIR-GECO variants, including dNIR-GECO0.2, dNIR-GECO0.6, dNIR-GECO0.8, and dNIR-GECO1.0 as purified proteins. Among the characterized variants, dNIR-GECO1.0 exhibited the strongest Ca^2+^-dependent response, with a Δ*F*/*F* of 22.1, representing a 20.1-fold improvement compared with the initial prototype dNIR-GECO0.2 (Δ*F*/*F* = 1.05), and a 2.8-fold and a 1.8-fold improvement over dNIR-GECO0.6 (Δ*F*/*F* = 8) and dNIR-GECO0.8 (Δ*F*/*F* = 12.1), respectively (**Supplementary Table 1**).

Spectroscopic characterization of dNIR-GECO1.0 revealed absorption peaks at 390 nm and 685 nm, which shifted to 392 nm and 679 nm upon Ca^2+^ binding (**Figure 2A**). The Ca^2+^-bound form showed excitation and emission peaks at 678 nm and 704 nm, respectively (**Figure 2B**). Ca^2+^ titration revealed an apparent dissociation constant (*K*_d,app_) of 181 nM and a Hill coefficient of 1.4, indicating enhanced cooperativity of Ca^2+^ binding compared with NIR-GECO2G (Hill coefficient, 0.78) (**Figure 2C**). pH titration of dNIR-GECO1.0 revealed a bell-shaped fluorescence profile, with apparent p*K*_a_ values of 5.00, 6.02 for the Ca^2+^-unbound form, and 5.05, 9.67 for the Ca^2+^-bound form, respectively (**Figure 2D, Supplementary Table 1**). The highest fluorescence intensity was observed at pH 6, while the maximum Δ*F*/*F* was observed at pH 9. Representative emission spectra across the measured pH range are provided in **Supplementary Figure 4**. dNIR-GECO1.0 exhibited an extinction coefficient of 68.4 mM^-1^cm^-1^, comparable to NIR-GECO2G in its bright state (74 mM^-1^cm^-1^), and a higher quantum yield (0.09 vs 0.06). As a result, dNIR-GECO1.0 has a molecular brightness 1.8-fold higher than NIR-GECO2G and 11.1-fold higher than dNIR-GECO0.2 (**Supplementary Table 1**). These results demonstrated that dNIR-GECO1.0 is a high-performance direct response NIR GECI with improved Ca^2+^ sensitivity and brightness.

**Figure 2.**
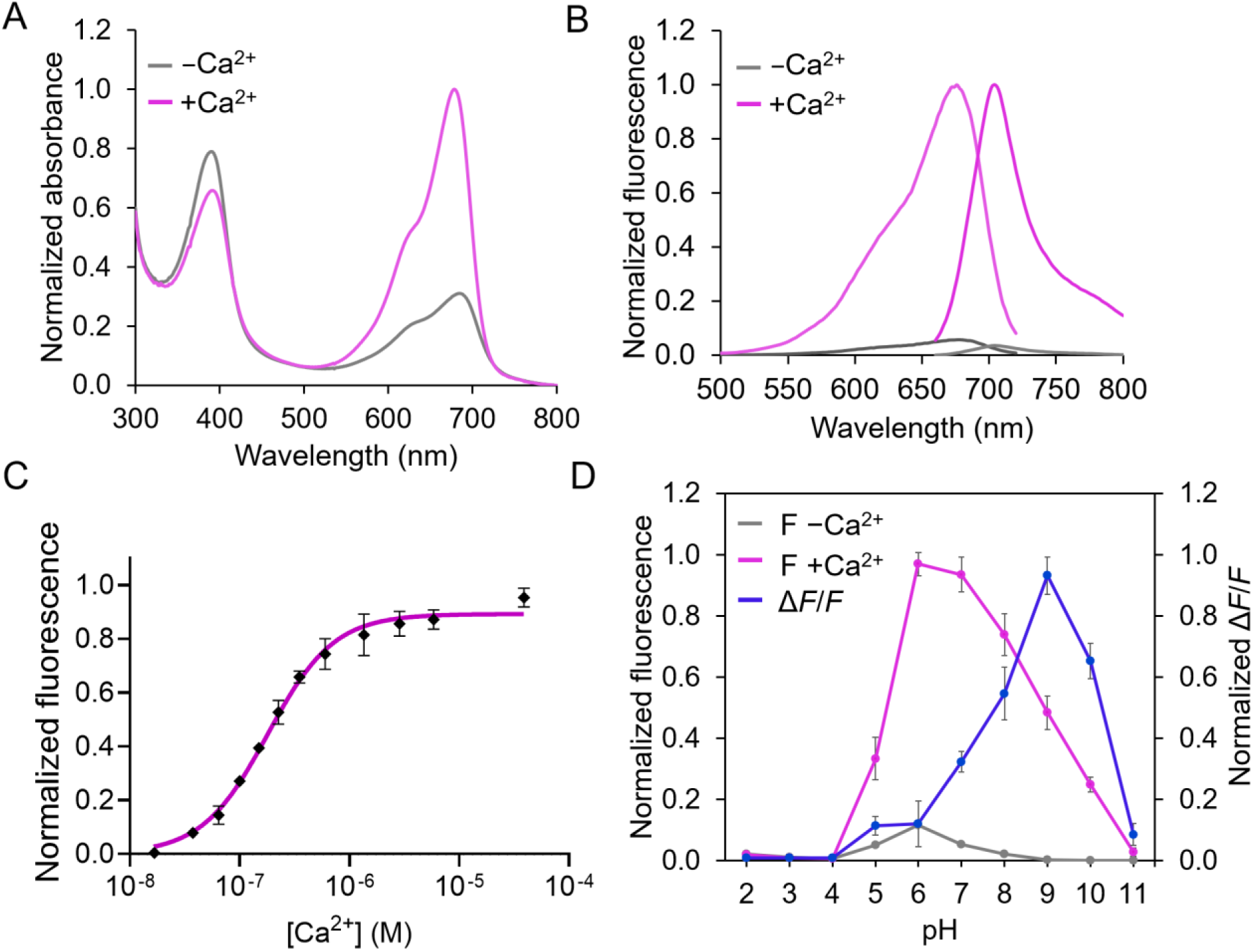
*In vitro* characterization of dNIR-GECO1.0. (A) Absorbance spectra of dNIR-GECO1.0 in the absence and presence of Ca^2+^. (B) Fluorescence excitation (emission at 770 nm) and emission spectra (excitation at 635 nm), with and without Ca^2+^. (C) Ca^2+^ titration curve of dNIR-GECO1.0 (mean ± SD, n = 3). (D) pH titration curves in the presence and absence of Ca^2+^ and Δ*F*/*F* measured at each pH (mean ± SD, n = 3).

### Characterization of dNIR-GECO variants in mammalian cells

To evaluate the performance of the engineered dNIR-GECO variants in mammalian cells, we characterized dNIR-GECO1.0, alongside dNIR-GECO0.8 and NIR-GECO2G, in HeLa cells. We first assessed the cellular brightness of these indicators in the resting state. Each indicator was co-expressed with EGFP as an internal reference via the self-cleaving peptide P2A, and the fluorescence was analyzed by flow cytometry in the presence or absence of 25 µM exogenous BV. Both dNIR-GECO variants exhibited detectable fluorescence without BV supplementation in HeLa cells, demonstrating their ability to incorporate endogenous BV. Supplementation with BV increased fluorescence by ∼6.3-fold for NIR-GECO2G, ∼17.1-fold for dNIR-GECO0.8, and ∼10.3-fold for dNIR-GECO1.0 (**Figure 3A**), suggesting that the BV availability limits the fluorescence brightness of these indicators in mammalian cells, with a major fraction of the expressed indicator remaining unbound to BV.

**Figure 3.**
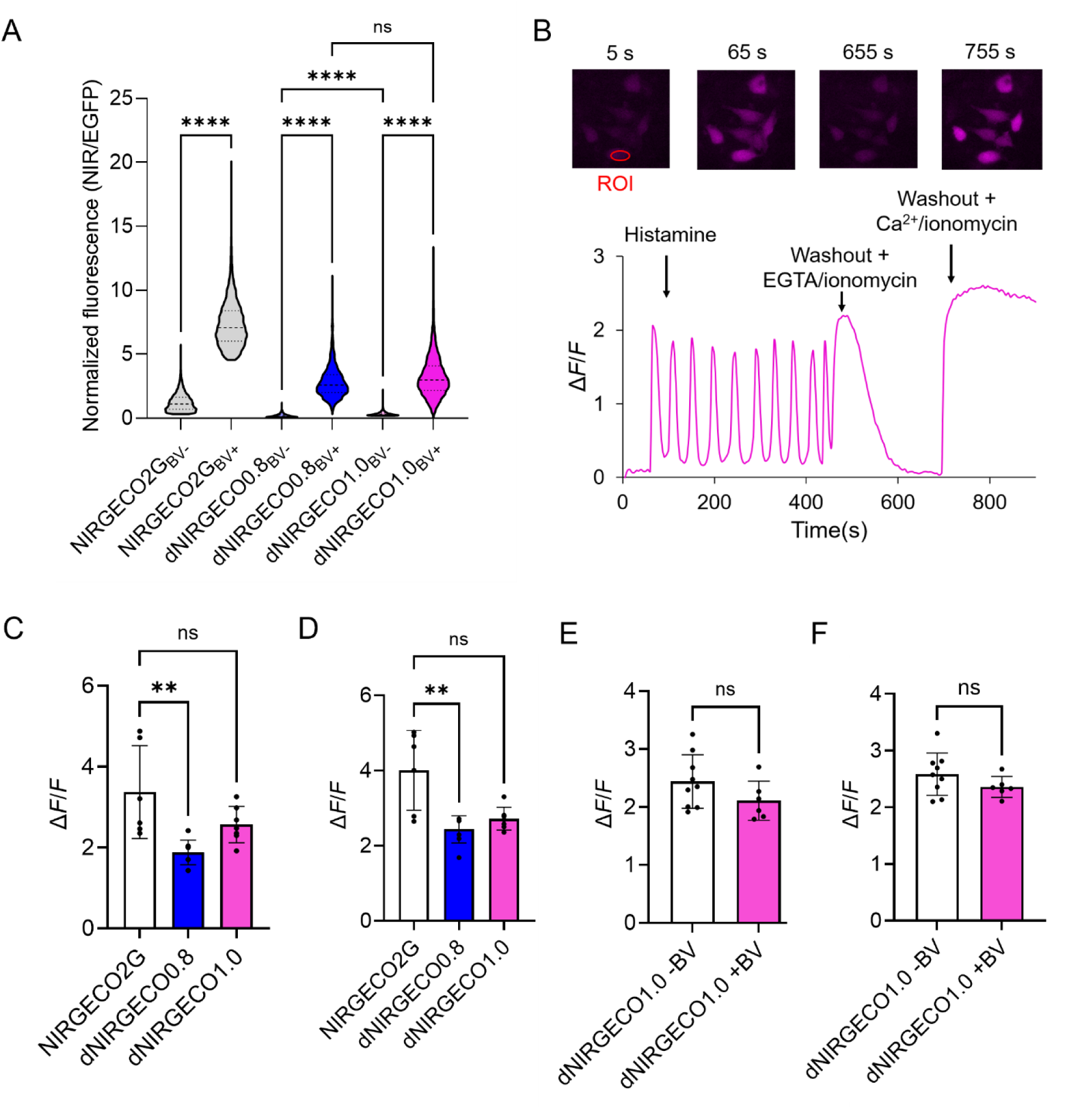
Characterization of dNIR-GECO variants and NIR-GECO2G in live mammalian cells. (A) Violin plots showing the basal fluorescence intensity of NIR-GECO2G, dNIR-GECO0.8, and dNIR-GECO1.0. NIR-GECO2G exhibits an inverse Ca^2+^ response, with higher fluorescence in the Ca^2+^-free state, whereas dNIR-GECO variants exhibit direct responses, with higher fluorescence in the Ca^2+^-bound state. Fluorescence intensity was measured by flow cytometry from approximately 1000-4000 transfected HeLa cells co-expressing P2A-EGFP in the presence or absence of 25 µM exogenous BV. n.s., not significant; ****p < 0.0001; Kruskal-Wallis test. (B) Representative fluorescence trace of a dNIR-GECO1.0 expressing HeLa cell without exogenous BV supplementation. Histamine was added to a final concentration of 10 µM at t = 1 min. Five minutes later, 5 mM EGTA/ionomycin was added to deplete Ca^2+^. Finally, 10 mM CaCl_2_/ionomycin was added to saturate the Ca^2+^ binding. (C) Histamine-induced Ca^2+^ responses of NIR-GECO2G, dNIR-GECO0.8, and dNIR-GECO1.0. The fluorescence response was calculated as |*F* − *F*₀|/*F*₀ for direct response indicators and |*F* − *F*₀|/*F* for inverse response indicators, where *F*₀ represents the resting fluorescence before histamine stimulation and *F* represents the peak fluorescence response (*F*_max_ for direct response indicators and *F*_min_ for inverse response indicators). This calculation was used to quantify the magnitude of the fluorescence change regardless of response direction. n = 6 cells. n.s., not significant; ****p < 0.0001; Kruskal-Wallis test (D) Ca_2+_-dependent fluorescence responses of NIR-GECO2G, dNIR-GECO0.8, and dNIR-GECO1.0. Δ*F*/*F* was calculated as (*F*_max_ − *F*_min_)/*F*_min_ for direct response indicators (dNIR-GECO variants) and (*F*_max_ − *F*_min_)/*F*_max_ for the inverse response indicator NIR-GECO2G. *F*_max_ and *F*_min_ represent the maximum and minimum fluorescence intensities under Ca^2+^-saturated and Ca^2+^-depleted conditions, respectively, for dNIR-GECO variants, and under Ca^2+^-depleted and Ca^2+^-saturated conditions, respectively, for NIR-GECO2G. n = 6 cells. n.s., not significant; ****p < 0.0001; Kruskal-Wallis test (E) Comparison of histamine-induced Δ*F*/*F* responses of dNIR-GECO1.0 in the absence and presence of exogenous BV. -BV, n = 9 cells; +BV, n = 6 cells. n.s., not significant; Mann-Whitney test. (F) Comparison of Ca^2+^/ionomycin-treated Δ*F*/*F* responses of dNIR-GECO1.0 in the absence and presence of exogenous BV. -BV, n = 9 cells; +BV, n = 6 cells. n.s., not significant; Mann-Whitney test.

To evaluate the Ca^2+^ response of dNIR-GECO1.0 in living cells, we treated HeLa cells expressing dNIR-GECO1.0 with 10 µM histamine, followed by sequential addition of ionomycin/EGTA and ionomycin/Ca^2+^. As expected, histamine treatment induced robust fluorescence oscillations, whereas subsequent addition of ionomycin/EGTA and ionomycin/Ca^2+^ caused a decrease and an increase in fluorescence, respectively (**Figure 3B**). We then quantified the maximum histamine-induced fluorescence changes for dNIR-GECO1.0 and compared it with that of dNIR-GECO0.8 and NIR-GECO2G. The maximum *ΔF*/*F* values were 2.7 ± 0.38 for dNIR-GECO1.0, 1.8 ± 0.34 for dNIR-GECO0.8, and 3.1 ± 1.14 for NIR-GECO2G (**Figure 3C**).

Similarly, the Δ*F*/*F* values in response to treatment with ionomycin/Ca^2+^ for dNIR-GECO1.0, dNIR-GECO0.8 and NIR-GECO2G were 2.7 ± 0.3, 2.4 ± 0.39, and 4.0 ± 1.1, respectively (**Figure 3D**). Addition of 25 µM BV improved the brightness of the cells but had no effect on the Δ*F*/*F* of dNIR-GECO1.0, indicating that the fraction in the BV unbound state does not affect the Ca^2+^-dependent response (**Figure 3E, F**).

### Identifying the mutations responsible for the direct response to Ca^2+^

In an attempt to identify key mutations responsible for the direct response of dNIR-GECO variants, we introduced each of the 11 mutations (5 mutations in the CaM domain, 1 mutation in the RS20 peptide and 5 mutations in the mIFP domain) from dNIR-GECO0.2 back into the inverse response NIR-GECO2G and tested their fluorescence responses from low to high Ca^2+^ levels in cell lysates (**Supplementary Table 4**). However, none of the individual mutations resulted in a direct response, suggesting that the direct response might not be attributed to a single mutation. We therefore shifted our focus from individual mutations to domain-level contributions. Using dNIR-GECO.8 as the template, we replaced the CaM domain, which has accumulated a large number of mutations during directed evolution, with the parental NIR-GECO2G CaM domain, and designated the resulting version as dNIR-GECO0.8 [N2 CaM]. Surprisingly, the CaM domain replacement reversed the response, converting direct response dNIR-GECO0.8 back into an inverse response indicator (**Supplementary Figure 5**), indicating that the mutations within CaM domain are responsible for the direct response of dNIR-GECO variants. To further confirm this finding, we tested 7 other dNIR-GECO variants found in the final rounds of directed evolution of dNIR-GECO1.0, including three variants that exhibited large Δ*F*/*F* but low brightness (Variant 1, Variant 2 and Variant 3) and four variants with both moderate Δ*F*/*F* and brightness (Variant 4, Variant 5, Variant 6 and Variant 7). Indeed, after swapping to the CaM domain of NIR-GECO2G, all of the 7 dNIR-GECO variants became reverse-response indicators (**Supplementary Figure 5** and **Supplementary Table 5**).

To identify the key mutations of CaM that flip the Ca^2+^-response type, we performed a bidirectional mutational analysis of the CaM domain in dNIR-GECO0.8 using cell lysates. Specifically, we examined whether introducing individual mutations into the inverse response variant dNIR-GECO0.8 [N2 CaM] could weaken or reverse its inverse response, and whether reverting these mutations in dNIR-GECO0.8 could diminish its direct response. We first introduced each of the 13 CaM mutations present in dNIR-GECO0.8 individually into dNIR-GECO0.8 [N2 CaM], generating 13 single-mutant variants. The resulting variants were then tested for their Ca^2+^-dependent fluorescence responses at lower (120 nM Ca^2+^) and high concentrations (39 µM and 0.5 mM Ca^2+^) (**Figure 4A**).

**Figure 4.**
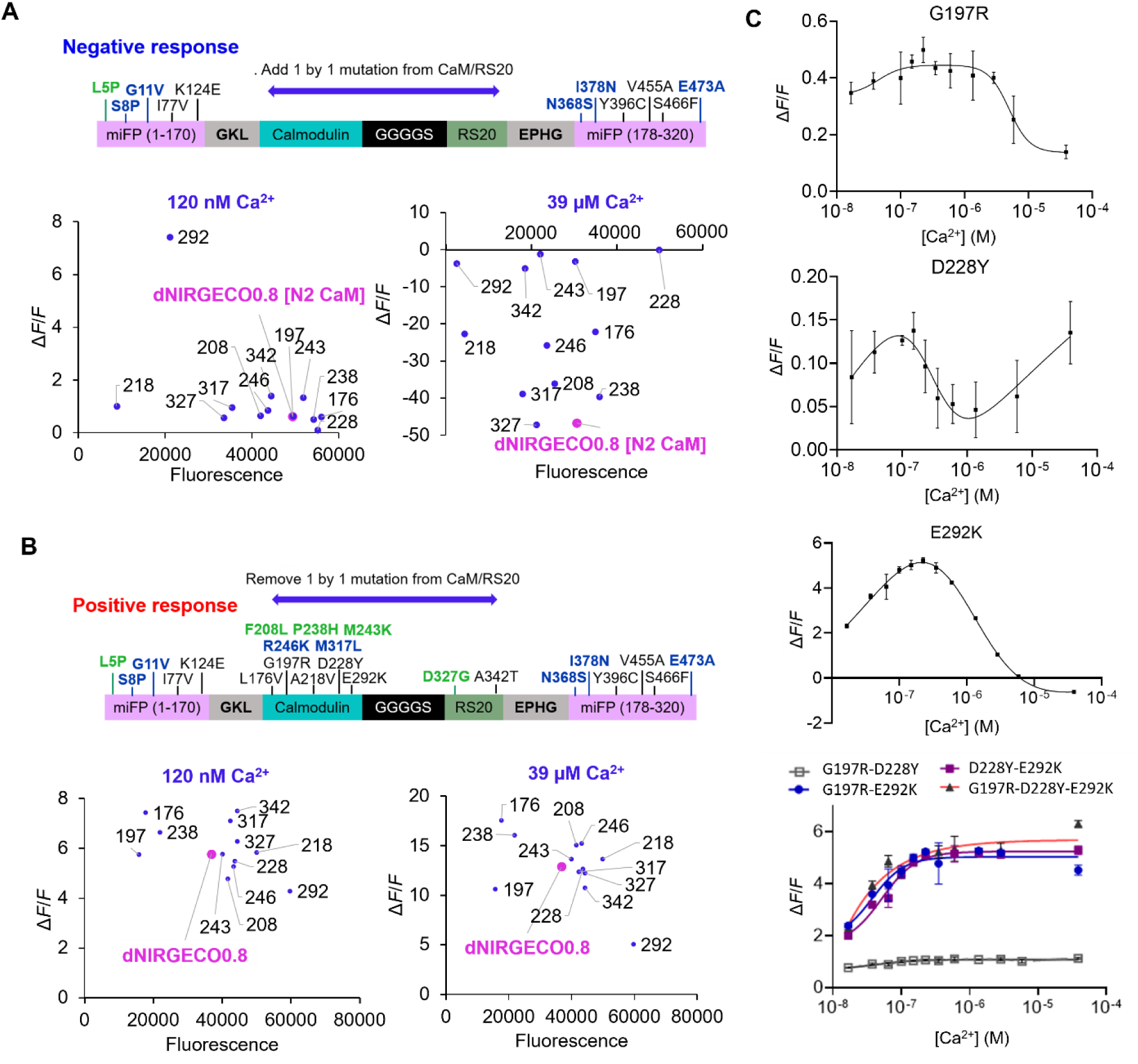
Identification of key CaM mutations responsible for the direct response in dNIR-GECO0.8. (A) Fluorescence changes of dNIR-GECO0.8 [N2 CaM] variants when Ca^2+^ concentration was increased from 0 nM to 120 nM (left) and 39 μM (right). dNIR-GECO0.8 [N2 CaM] variants were generated by introducing one of the 13 mutations accumulated on the CaM domain of dNIR-GECO0.8. (B) Fluorescence changes of dNIR-GECO0.8 variants when Ca^2+^ concentration was increased from 0 nM to 120 nM (left) and 39 μM (right). dNIR-GECO0.8 mutations were generated by reverting one of the 13 mutations accumulated on the CaM domain of dNIR-GECO0.8 (C) Ca^2+^ titration curves of dNIR-GECO0.8 [N2 CaM] variants containing mutations critical for the direct response switching (mean ± SD, n = 3).

This experiment revealed 5 mutations that significantly weakened or diminished the inverse response, namely Gly197Arg, Asp228Tyr, Met243Lys, Glu292Lys, and Ala342Thr. Among the 5 mutations, Glu292Lys produced a strong direct response at 120 nM Ca^2+^ (Δ*F*/*F* = 7.4), but the response switched back to an inverse type at a higher Ca^2+^ concentration of 39 µM (Δ*F*/*F* = −3.7) (**Figure 4A and Supplementary Table 6**). We next individually reverted the same 13 mutations in the direct response variant dNIR-GECO0.8 to evaluate their contributions to its direct fluorescence response. Although no single mutation completely abolished the direct response phenotype, reversion of Glu292Lys resulted in a substantial reduction in response amplitude, with Δ*F*/*F* decreasing from 12.9 to 5.1 at 39 µM Ca^2+^, corresponding to a 60.5% decrease (**Figure 4B, Supplementary Table 7**). Together, these results indicate that Glu292Lys plays a major role in enhancing and stabilizing the direct response behavior of dNIR-GECO0.8.

Having identified 5 candidate mutations contributing to the switch from inverse to direct response, with Glu292Lys emerging as a major contributor, we next investigated their combined effects. We introduced all 10 possible pairwise combinations of these mutations into dNIR-GECO0.8 [N2 CaM] and characterized their Ca^2+^-responses (**Supplementary Table 8**). Interestingly, all combinations containing Glu292Lys exhibited strong direct responses. Among them, the Asp228Tyr+Glu292Lys and Gly197Arg+Glu292Lys variants showed direct responses at both low and high Ca^2+^ concentrations, suggesting that these mutation combinations are particularly important for establishing the direct response behavior. Combining all three mutations (Gly197Arg, Asp228Tyr, and Glu292Lys) further improved the direct response, yielding a variant termed dNIR-GECO0.8 [N2 CaM] 3M.

To better characterize these variants, Ca^2+^ titration assays were performed using purified proteins. The variants containing individual mutations exhibited biphasic response curves, whereas the double-mutant variants (Gly197Arg+Asp228Tyr, Asp228Tyr+Glu292Lys and Gly197Arg+Glu292Lys) and the triple-mutant variant (Gly197Arg+Asp228Tyr+Glu292Lys) displayed sigmoidal direct response curves against Ca^2+^ (**Figure 3C**). These results revealed that the combination of Gly197Arg, Asp228Tyr, and Glu292Lys are necessary to convert the biphasic response into a stable direct response biosensor. To further confirm this hypothesis, we introduced the three key mutations (Gly197Arg, Asp228Tyr, and Glu292Lys) into the parental NIR-GECO2G instead of dNIR-GECO0.8 [N2 CaM], generating the NIR-GECO2G3M variant. NIR-GECO2G3M showed a small but apparent direct response (Δ*F*/*F* = 0.12) (**Supplementary Figure 6A, B**), confirming that the three mutations are the minimal set required to convert NIR-GECO2G into a direct response Ca^2+^ indicator. These three mutations had been initially introduced in the first two rounds of evolution for a direct response NIR GECI prototype (**Figure 1B**).

### Resonance Raman spectroscopy of NIR-GECO2G and dNIR-GECOs

To investigate the structural basis of direct response vs inverse response NIR-GECO variants, we performed resonance Raman (RR) spectroscopy to examine the BV chromophore (**Figure 5A**). We measured RR spectra of NIR-GECO2G and the two dNIR-GECO variants, dNIR-GECO1.0 and dNIR-GECO0.8 [N2 CaM] 3M with excitation at 405 nm, which is resonant with the absorption band at 390 nm (**Figure 5B**). For the spectra in **Figure 5B**, the top trace represents the Ca^2+^-free protein, and the bottom represents the Ca^2+^-bound protein. Since NIR-GECO2G exhibits an inverse fluorescence response to Ca^2+^-binding, while the dNIR-GECOs exhibit a direct fluorescence response, the red traces correspond to spectra in the brighter state, and the blue traces are those in the dimmer state. For all three indicators, the spectral shapes in the bright state and the dimmer state exhibited similar characteristics. Specifically, in response to Ca^2+^-binding to the proteins, bands shifted or appeared around 1600 cm^-1^, 1247 cm^-1^, and 810 cm^-1^. To clearly highlight these spectral changes, we derived the difference spectra in response to Ca^2+^-binding (**Supplementary Figure 7**). The difference spectrum upon Ca^2+^-binding in NIR-GECO2G has a mirror relationship with the difference spectra of the dNIR-GECOs, consistent with the reversal in the direction of the change in the fluorescence response. We conclude that identical structural changes are occurring in the BV chromophore, but in opposite directions.

**Figure 5.**
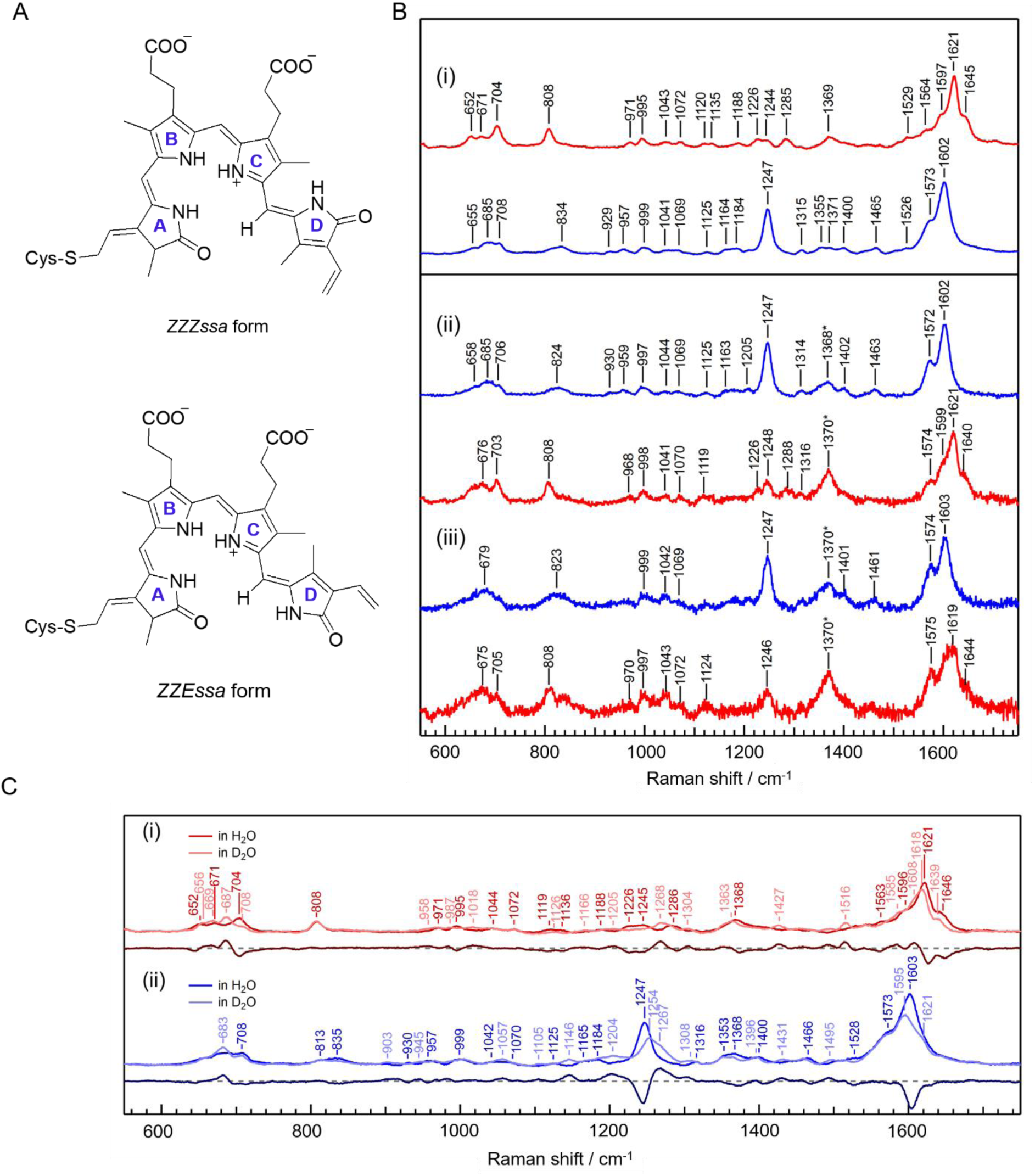
Resonance Raman (RR) spectroscopic analysis of NIR-GECOs. (A) Chemical structures of biliverdin (BV) in the *ZZZ* and *ZZE* configurations. (B) RR spectra of (i) NIR-GECO2G, (ii) dNIR-GECO1.0, and (iii) dNIR-GECO0.8 [N2 CaM] 3M, excited at 405 nm. For each panel, upper and lower traces represent the spectra of the Ca^2+^-free and Ca^2+^-bound proteins, respectively. The red traces display the spectra in the bright fluorescent state, and the blue ones exhibit those in the dimmer fluorescent state. The fluorescence background has been subtracted from each spectrum. The bands at ∼1370 cm^−1^ appearing in the dNIR-GECOs, which are marked with asterisks, are due to the impurity bands that could not be completely removed during the purification process. (C) Deuteration effects on the RR spectra of (i) Ca^2+^-free and (ii) Ca^2+^-bound NIR-GECO2G. For each panel, the upper two traces are the RR spectra in the H_2_O (light color) and D_2_O (dark color) buffers. Lower traces represent the difference between the spectra measured in the H_2_O and D_2_O buffers. The fluorescence background has been subtracted from each spectrum.

For NIR-GECO2G, we examined the effect of deuteration on the RR bands (**Figure 5C**). In D_2_O, the proton on the pyrrole NH group is replaced with a deuterium. As a result, frequency shifts are observed in the bands due to the vibrational modes coupled to the vibrational mode of the pyrrole NH/D group. By comparing the spectral features of the BV chromophore in NIR-GECOs, the spectral changes induced by the H/D exchange, and the reported RR spectra of the Pr and Pfr states of phytochromes containing the BV chromophore^22,23^, we offer the following interpretation of the observed RR bands.

The bands appearing at 1646, 1621, and 1596 cm^-1^ in the bright state and 1603, 1573 cm^-1^ in the dimmer state are assigned to the C=C stretching modes of the methine bridge^22,23^. Upon deuteration, the bands at 1646, 1621, and 1596 cm^-1^ shifted to 1639, 1618, and 1585 cm^-1^ in the bright state, and the 1603 cm^-1^ band in the dimmer state shifted to 1595 cm^-1^ in the dimmer state. These spectral features were consistent with the the Pr and Pfr states of phytochrome^22,23^. The C=C stretching bands serve as structural markers that determine the configuration of the BV chromophore, particularly the orientation of ring D. Consequently, we propose that the BV chromophores in the bright and dimmer states of NIR-GECOs adopt the *ZZZssa* and *ZZEssa* configurations, respectively.

The strong bands at 1621 cm^-1^ in the bright state and at 1603 cm^-1^ in the dimmer state are attributed to the C=C stretching mode in the methine bridge between the C–D rings^22,23^. As the distortion along the C–D bridge increases, the degree of π-electron conjugation weakens and the double-bond character increases, resulting in the band appearing at higher frequency. Therefore, we speculate that the distortion of the C–D bridge in the bright state is larger than that in the dimmer state. The observation was consistent with the appearance of the hydrogen out-of-plane wagging band in the methine bridge between the C–D rings (808 cm^-1^ for the bright state)^22,23^, which is observed in the spectra of the distorted polyene chain. The bands appearing at 1563 cm^-1^ in the bright state and 1528 cm^-1^ in the dimmer state are attributed to NH rocking modes^22,23^, and in D_2_O these ND rocking bands shifted to 1018 and 1057 cm^-1^, respectively. The NH rocking frequency increases as hydrogen bonds at the pyrrole NH groups become stronger. Therefore, the fact that the NH rocking band appears at a higher frequency in the bright state suggests that strong hydrogen bonds are formed at the pyrrole rings.

The strong 1247 cm^-1^ band in the dimmer-state spectrum was shifted upon the H/D exchange, and is attributed to the vibrational mode coupled between the C–C/C–N stretching and CH rocking modes and the NH rocking modes^22,23^. Structural changes during the electronic transition between the ground and excited states give rise to the *A*-term RR enhancement arising from the Franck-Condon mechanism^24^. A similar band has been observed in the 405-nm excited RR spectra of the BV chromophore of sandercyanin^25^. In sandercyanin, the strong 1247 cm^-1^ band was correlated with increased space around the D ring and, presumably, greater structural flexibility of the BV chromophore.

## DISCUSSION

Developing far-red and NIR GECIs with performance comparable to that of widely used green and red fluorescent GECIs has been a long-standing goal for biosensor engineers. However, existing NIR GECIs lag behind their visible-wavelength counterparts in several key aspects, including undesirable fluorescence response type (i.e. inverse response), low brightness, and poor photostability. Here, we addressed one of these major limitations by developing dNIR-GECO1.0, a direct response NIR calcium indicator engineered from the inverse response NIR-GECO2G scaffold, through multiple rounds of directed evolution and protein engineering.

As the newest member of the phytochrome-based far-red and NIR fluorescent Ca^2+^ indicator family, dNIR-GECO1.0 provides several advantages over previously developed sensors. Unlike the earlier NIR-GECO series (NIR-GECO1^14^, NIR-GECO2^15^, NIR-GECO2G^15^, and NIR-GECO3^18^), which exhibit inverse fluorescence changes upon Ca^2+^ binding, the fluorescence of dNIR-GECO1.0 increases upon Ca^2+^ binding, providing a robust readout of Ca^2+^ dynamics. Compared with chemigenetic NIR GECIs such as HaloCaMP and WHaloCaMP^20,21^, which require exogenous ligand supplementation for fluorescence readout, dNIR-GECO1.0 is fully genetically encoded and can be directly imaged in mammalian cells. Importantly, dNIR-GECO1.0 showed a large Ca^2+^-dependent fluorescence response in purified protein (Δ*F*/*F* = 22.1) while maintaining favorable photophysical properties, including high molecular brightness (18.1 mM^-1^ cm^-1^) and near-infrared excitation and emission. The dynamic range of dNIR-GECO1.0 is substantially larger than the previously reported direct response phytochrome-based indicators, including GAF-CaMP2-sfGFP^11^ and GAF-CaMP3-sfGFP^12^, which also have lower molecular brightness (3.7 and 2.9 mM^-1^ cm^-1^ respectively) and required sfGFP fusion and exogenous biliverdin supplementation to achieve detectable fluorescence in mammalian cells. In summary, dNIR-GECO1.0 enables detection of Ca^2+^ dynamics in mammalian cells in the near-infrared window without external chemical addition, and without the drawbacks and complications associated with inverse response NIR GECIs.

The development of dNIR-GECO1.0 also highlights the importance of balancing dynamic range and cellular brightness of a biosensor during engineering. We found several intermediate dNIR-GECO variants that showed exceptionally large Ca^2+^ responses in purified proteins but were not selected as the final variant due to their dim fluorescence in HeLa cells. These results indicate that improvements measured in purified proteins do not always translate directly to cellular performance, likely due to differences in BV incorporation and protein stability between bacteria and mammalian cells. Using a previously established dual screening strategy^14^, we combined purified protein screening with mammalian cell-based screening to select variants that exhibit both large Ca^2+^ responses in purified proteins and bright cellular fluorescence in HeLa cells. This strategy has been proven effective and ultimately led to the discovery of the final dNIR-GECO variant, although there remains a room to improve its cellular brightness as indicated by the BV-addition experiment.

To understand the molecular basis of the response type switching, we investigated the mutations accumulated during dNIR-GECO evolution. and identified three key mutations, Gly197Arg, Asp228Tyr, and Glu292Lys, which, in combination, were necessary to convert the inverse response NIR-GECO2G scaffold into a direct response Ca^2+^ indicator. These findings demonstrate that fluorescence response direction can be reprogrammed through protein engineering. A similar finding was also observed during the development of eLACCO1, in which linker optimization followed by directed evolution successfully changed the response from inverse to direct response^26^.

RR spectroscopy analysis revealed that the structural changes in the BV chromophore between the bright and dimmer fluorescent states of NIR-GECO series are identical regardless of the direction of the fluorescent response to Ca^2+^ binding. In the bright fluorescent state, the BV chromophore is in the *ZZZssa* configuration, in which the methine group between the C and D rings adopts a twist and the pyrrole ring NH group forms strong hydrogen bonds, resulting in a rigid structure. In contrast, the chromophore in the dimmer state adopts a *ZZEssa* configuration and exhibits a flexible structure with space near the chromophore.

Overall, our study demonstrates that the fluorescence response direction of a genetically encoded Ca^2+^ indicator can be reprogrammed through directed evolution. The resulting dNIR-GECO1.0 is a robust tool to report Ca^2+^ dynamics in mammalian cells. Future endeavors could focus on further improving its performance in neurons and *in vivo*.

## MATERIALS AND METHODS

### Preparation of NIR-GECO variants

Genes encoding NIR-GECO2G and dNIR-GECO variants were cloned into the pcDuEx2 vector, which co-expresses heme oxygenase-1 (HO-1) from the cyanobacterium *Synechocystis* to enable BV production in *E. coli*. This expression system was used for both *in vitro* protein characterization and mammalian cell imaging. For large-scale protein purification, plasmids were instead expressed using a pBAD vector containing HO-1 to enhance BV production.

PCR amplification was performed using Phusion High-Fidelity DNA Polymerase (Thermo Fisher Scientific). Site-directed mutagenesis was conducted using the revised QuikChange mutagenesis method. Plasmid DNA was purified using the GeneJET Plasmid Miniprep Kit (Thermo Fisher Scientific) according to the manufacturer’s instructions.

### Engineering and development of dNIR-GECO variants

NIR-GECO2G was used as the template for directed evolution. Error-prone PCR was performed using Taq DNA polymerase (New England Biolabs) and optimized by adjusting the manganese concentration to achieve the desired mutation rate per round. The amplified gene fragments were purified using the GeneJET Gel Extraction Kit (Thermo Fisher Scientific) and ligated into the pDuex2 vector using XhoI and HindIII restriction enzymes (Thermo Fisher Scientific).

The ligation products were transformed into ElectroMAX DH10B competent *Escherichia coli* cells (Invitrogen). Transformants were plated on 12–20 agar plates containing ampicillin and incubated overnight at 37 °C. For each round of screening, approximately 200–300 colonies were screened per plate across 12–20 plates, and the brightest 10% of colonies were selected. Selected colonies were cultured in 1 mL LB medium supplemented with 100 µg mL^-1^ ampicillin in 96-well deep-well plates.

Cultures were incubated overnight at 37 °C with shaking at 250 rpm, followed by additional incubation overnight at room temperature. The following day, cells were collected by centrifugation, and the supernatants were discarded. Cell pellets were resuspended in 100 µL B-PER reagent (Thermo Fisher Scientific), vortexed gently for 20 min, and centrifuged at 4,000 × g for 5 min. The resulting supernatants were used for fluorescence screening.

Fluorescence measurements were performed in 96-well plates using a Tecan microplate reader. The +Ca^2+^ buffer contained 30 mM MOPS, 100 mM KCl, and 10 mM CaEGTA (pH 7.2), whereas the -Ca^2+^ buffer contained 30 mM MOPS, 100 mM KCl, and 10 mM EGTA (pH 7.2). Fluorescence was measured using excitation at 630 nm and emission scanning from 675-735 nm with 2 nm increments. Promising variants were identified, sequenced, and used as templates for subsequent rounds of directed evolution.

### Protein purification and *in vitro* characterization of dNIR-GECO proteins

NIR-GECO2G and dNIR-GECO variants were expressed in DH10B *E. coli* cells containing pBAD-HO1 plasmids. Overnight starter cultures were prepared in 10 mL Terrific Broth (TB) supplemented with 100 µg mL^-1^ ampicillin and incubated at 37 °C with shaking at 250 rpm.

Cultures were transferred into 100 mL TB medium containing ampicillin and grown at 37 °C until the optical density at 600 nm (OD600) reached 0.6 - 1.0. Protein expression was induced by adding 0.2% (w/v) L-arabinose, followed by overnight incubation at room temperature with shaking at 250 rpm. Cell pellets were lysed in a lysis buffer containing 10 mM imidazole, 20 mM Na_2_HPO_4_, and 300 mM NaCl (pH 7.2) using a Branson cell disruptor with 2 min pulses separated by 1 min intervals. Lysates were centrifuged at 26,953 × g for 45 min, and supernatants were filtered through 0.22 µm Millex-GV filters (Merck Millipore). Filtered lysates were loaded onto 10 mL Pierce centrifuge columns (Thermo Fisher Scientific) containing pre-equilibrated nickel-nitrilotriacetic acid (Ni-NTA) resin (G-Biosciences). Samples were incubated on a mechanical rotor for 1 h at 4 °C and washed several times with washing buffer containing 20 mM imidazole, 20 mM Na_2_HPO_4_, and 300 mM NaCl (pH 7.2). Proteins were eluted using buffer containing 250 mM imidazole, 20 mM Na_2_HPO_4_, and 300 mM NaCl (pH 7.2), and concentrated using Amicon Ultra-0.5 centrifugal filters (Merck Millipore). Concentrated proteins were subjected to multiple buffer exchange steps into 30 mM MOPS and 100 mM KCl buffer (pH 7.2). Fluorescence measurements were performed using the same procedure as described for B-PER lysates. For Raman analysis, proteins were expressed in 5 L cultures and purified by Ni-NTA affinity chromatography as described above, with modifications to the cell lysis and purification procedure for the larger culture volume. The purified proteins were then buffer-exchanged into 30 mM MOPS and 100 mM KCl buffer (pH 7.2), followed by size-exclusion chromatography using a Superdex 200 Increase 10/300 GL column equilibrated with SEC buffer (20 mM HEPES-NaOH (pH 7.4) and 150 mM NaCl). The peak fractions were collected for further analysis.

### Characterization of purified proteins

Absorption spectra under Ca^2+^-free and Ca^2+^-saturated conditions were measured using a Lambda 950 spectrophotometer (PerkinElmer) in 1 cm path-length quartz cuvettes over a wavelength range of 300-800 nm. Fluorescence spectra were recorded using a Tecan microplate reader over a wavelength range of 500–800 nm. Extinction coefficients were estimated by comparing absorbance at 678 nm with absorbance at 391 nm, using an extinction coefficient of 39,900 M^-1^ cm^-1^ at 391 nm. Absolute quantum yields were measured using an integrating sphere spectrometer (Quantaurus-QY, Hamamatsu). Ca^2+^ titration experiments were performed to determine apparent Ca^2+^-binding affinity. Buffers containing free Ca^2+^ concentrations ranging from 0 µM to 39 µM were prepared by mixing 10 mM EGTA and 10 mM CaEGTA solutions at pH 7.2. Apparent dissociation constants (*K*_d,app_) and Hill coefficients were determined by sigmoidal fitting using Prism software (GraphPad Software).

Apparent p*K*_a_ values were determined by diluting purified proteins into buffers containing 30 mM trisodium citrate, 30 mM sodium borate, 30 mM MOPS, 100 mM KCl, and either 5 mM CaCl_2_ or 5 mM EGTA, adjusted to pH values ranging from 2-11. The p*K*_a_ values were determined from the inflection points of sigmoidal fits of fluorescence intensity versus pH.

### Characterization in HeLa cells

HeLa cells were cultured in Dulbecco’s Modified Eagle Medium (DMEM, Gibo) containing 4.5g/L glucose and supplemented with 10% fetal bovine serum (FBS; Sigma-Aldrich) and 1% penicillin–streptomycin (Nacalai Tesque). Cells were maintained at 37 °C in a humidified incubator containing 5% CO_2_. Cells were seeded onto 35 mm glass-bottom culture dishes (Iwaki) and transfected with 3 µg plasmid DNA using polyethyleneimine (PEI) in Opti-MEM Reduced Serum Medium (Thermo Fisher Scientific). The culture medium was replaced 4 h after transfection. Forty-eight hours after transfection, cells were washed twice with phenol red-free Hank’s Balanced Salt Solution (HBSS) containing Ca^2+^ and Mg^2+^ and supplemented with 10 mM HEPES (pH 7.2). For biliverdin-supplemented conditions, cells were incubated with 25 µM biliverdin for 2 h prior to imaging.

At 48-72 h after transfection, live-cell fluorescence imaging was performed using an IX83 wide-field fluorescence microscope (Olympus) equipped with a pE-300 White LED light source (CoolLED), a 40× oil-immersion objective lens (NA = 1.3), an ImagEM X2 EM-CCD camera (Hamamatsu), and CellSens software (Olympus). Fluorescence images were acquired using a Cy5.5 filter set (675 nm excitation and 694 nm emission). Image analysis was performed using ImageJ software (National Institutes of Health).

For characterization of Ca^2+^-dependent fluorescence responses, cells were imaged for 1 min before stimulation with 10 µM histamine. After histamine washout, 5 mM EGTA was added to chelate extracellular Ca^2+^ and obtain the Ca^2+^-free state. Subsequently, 10 mM CaCl_2_ was added to achieve the Ca^2+^-saturated state. Image analysis was performed using ImageJ, and regions of interest (ROIs) were manually selected for individual cells.

For flow cytometry analysis, cells were transfected with 2 µg plasmids encoding pcDNA3.1-NIR-GECO2G-P2A-EGFP, pcDNA3.1-dNIRGECO0.8-P2A-EGFP, or pcDNA3.1-dNIRGECO1.0-P2A-EGFP and incubated for at least 48 h before analysis. HeLa cells expressing the variants were dissociated using trypsin and collected by centrifugation at 2,000 g for 2 min. For biliverdin-supplemented samples, cells were incubated with 25 µM biliverdin for 2 h prior to analysis. Cells were resuspended in HBSS supplemented with 10 mM HEPES and analyzed using sh800 (SONY). 488 and 638 nm lasers were used for excitation, and 525/50 and 720/60 bandpass filters were used to detect emission from EGFP and NIRGECO variants, respectively. Compensation was performed using cells expressing pcDNA3.1-EGFP and pcDNA3.1-NIRGECO2G as single-color positive controls and non-transfected cells for negative control. Transfected cells were gated based on the EGFP signal and the cellular brightness was calculated as the ratio of the NIRGECO variant signal to the EGFP signal.

### Resonance Raman (RR) measurements

A 405-nm diode laser (Coherent, SureLock LM-405) was used as the excitation light source for resonance Raman (RR) scattering. The excitation light was directed onto the sample solution in a φ10 NMR tube as a spinning cell. Backscattered light at 135 degrees was directed into the spectrometer system using a pair of achromatic lenses. Rayleigh scattered light was removed using a prism spectrometer (Bunkoukeiki, PF-200) as a prefilter, and the dispersed Raman scattered light was detected by a liquid nitrogen-cooled CCD camera (Teledyne, PyLon-2KBUV) using a spectrometer (HORIBA Jobin Yvon, iHR550). The excitation power at the sample position was 1 mW to avoid detection of unwanted photoreacted products in the observed spectra. The Raman shift was calibrated based on the Raman bands of cyclohexane, toluene, and acetone, with an error within ±1 cm^-1^. All measurements were performed at room temperature.

## AUTHOR CONTRIBUTIONS

M.Musa, R.H., and T.K. performed directed evolution and contributed to the development of dNIR-GECO. M.Musa carried out the final optimization and characterization. Y.Q. contributed to the development of the initial direct response prototype. M.Mizuno performed and analyzed Raman experiments and prepared the corresponding text. M.Musa, K.T., and R.U. performed large-scale Ni-NTA affinity protein purification for Raman analysis. K.S. performed further protein purification using size-exclusion chromatography under the supervision of O.N. S.I. performed FACS analysis. Y.N., T.T., Y.Q., and R.E.C. supervised the work. M.Musa drafted the original manuscript based on experimental findings and theses by R.H. and T.K, while Y.Q. and R.E.C. contributed extensively to writing and revising the manuscript. All authors reviewed and approved the final manuscript.

## ACKNOWLEDGEMENTS

Work in the lab of R.E.C. was supported by grants from the Japan Society for the Promotion of Science (JSPS) KAKENHI (24H00489 and 24H02267), JST CREST (JPMJCR25T3), and the National Institutes of Health (NIH) (U01NS123658). T.T. was supported by KAKENHI (23H02101, 26K01641, and 26H01674) and the Precise Measurement Technology Promotion Foundation. Y.N. was supported by JSPS KAKENHI (24K01621 and 25K01642) and JST (JPMJPR22E9 and JPMJCR24T6). Financial support of M.Mizuno is provided by KAKENHI (24H02263 and 24K01436). S.I was supported by JST ACT-X (JPMJAX2532). K.S was supported by JSPS KAKENHI (JP25KJ0866). O.N was supported by the Japan Agency for Medical Research and Development (AMED) (JP223fa627001) and JSPS KAKENHI (26H00007).

## DATA AVAILABILITY STATEMENT

All data supporting the findings of this study are included in the article and its Supplementary Information. Any additional data are available from the corresponding author upon reasonable request.

