## Supplementary Information for "A near-infrared fluorescent genetically encoded calcium ion biosensor with a direct response"

### Supplementary Figures

```
NIRGECO2G     MSVPLTTSAFGHAFLANCEREQIHLAGSIQPHGILLAVKEPDNVVIQASINAAEFLNTNFVVGRP  65
dNIRGECO0.2   MSVPLTTSAFGHAFLANCEREQIHLAGSIQPHGILLAVKEPDNVVIQASINAAEFLNTNFVVGRP
dNIRGECO0.4   MSVPLTTSAFGHAFLANCEREQIHLAGSIQPHGILLAVKEPDNVVIQASINAAEFLNTNFVVGRP
dNIRGECO0.6   MSVPLTTPAFVHAFLANCEREQIHLAGSIQPHGILLAVKEPDNVVIQASINAAEFLNTNFVVGRP
dNIRGECO0.8   MSVPPTTPAFVHAFLANCEREQIHLAGSIQPHGILLAVKEPDNVVIQASINAAEFLNTNFVVGRP
dNIRGECO1.0   MSVPPTTPAFVHAFLANCEREQIHLAGSIQPHGILLAVKEPDNVVIQASINAAEFLNTNFVVGRP

NIRGECO2G     LRDLGGDLPLQILPHLNGPLHLAPMTLRCTVGSPPRRVDCTIHRPSNGGLIVELEPATKATNIAP  130
dNIRGECO0.2   LRDLGGDLPLQVLPHLNGPLHLAPMTLRCTVGSPPRRVDCTIHRPSNGGLIVELEPATEATNIAP
dNIRGECO0.4   LRDLGGDLPLQVLPHLNGPLHLAPMTLRCTVGSPPRRVDCTIHRPSNGGLIVELEPATEATNIAP
dNIRGECO0.6   LRDLGGDLPLQVLPHLNGPLHLAPMTLRCTVGSPPRRVDCTIHRPSNGGLIVELEPATEATNIAP
dNIRGECO0.8   LRDLGGDLPLQVLPHLNGPLHLAPMTLRCTVGSPPRRVDCTIHRPSNGGLIVELEPATEATNIAP
dNIRGECO1.0   LRDLGGDLPLQVLPHLNGPLHLAPMTLRCTVGSPPRRVDCTIHRPSNGGLIVELEPATEATNIAP

NIRGECO2G     ALVGALHRITSSSSLMGLCDETATIFREITGFDRVMVMRLGALDDLTEEQIAEIKEAFSLFDKDG  195
dNIRGECO0.2   ALVGALHRITSSSSLMGLCDETATIFREITGFDRVMVMRLGALDDVTEEQIAEIKEAFSLFDKDG
dNIRGECO0.4   ALVGALHRITSSSSLMGLCDETATIFREITGFDRVMVMRLGKLDDVTEEQIAEIKEAFSLFDKDG
dNIRGECO0.6   ALVGALHRITSSSSLMGLCDETATIFREITGFDRVMVMRLGKLDDVTEEQIAEIKEAFSLFDKDG
dNIRGECO0.8   ALVGALHRITSSSSLMGLCDETATIFREITGFDRVMVMRLGKLDDVTEEQIAEIKEAFSLFDKDG
dNIRGECO1.0   ALVGALHRITSSSSLMGLCDETATIFREITGFDRVMVMRLGKLDDVTEEQIAEIKEAFSLFDKDG

NIRGECO2G     DGTITTKELGTVFRSLGQNPTEAELQDMINEVDADGDGIFDFPEFLTMMARKMNDSDSEEEIRGA  260
dNIRGECO0.2   DRTITTKELGTVFRSLGQNPTEVELQDMINEVYADGDGIFDFPEFLTMMARKMNDSDSEEEIRGA
dNIRGECO0.4   DRTITTKELGTVFRSLGQNPTEVELQDMINEVYADGDGIFDFPEFLTMMARKMNDSDSEEEIRGA
dNIRGECO0.6   DRTITTKELGTVFRSLGQNPTEVELQDMINEVYADGDGIFDFPEFLTMMAKKMNDSDSEEEIRGA
dNIRGECO0.8   DRTITTKELGTVLRSLGQNPTEVELQDMINEVYADGDGIFDFHEFLTKMAKKMNDSDSEEEIRGA
dNIRGECO1.0   DRTITTKELGTVLRSLGQNPTEAELQDMINEVYADGDGIFDFHEFLTKMAKKMNDSDSEEEIRGA

NIRGECO2G     FRVFDKDGNGYIGAAELRHVMTDLGEKLTDEEVDEMIRVADNDGDGQVNYEEFVQMMTAKGGGGS  325
dNIRGECO0.2   FRVFDKDGNGYIGAAELRHVMTDLGEKLTDEKVDEMIRVADNDGDGQVNYEEFVQMMTAKGGGGS
dNIRGECO0.4   FRVFDKDGNGYIGAAELRHVMTDLGEKLTDEKVDEMIRVADNDGDGQVNYEEFVQMMTAKGGGGS
dNIRGECO0.6   FRVFDKDGNGYIGAAELRHVMTDLGEKLTDEKVDEMIRVADNDGDGQVNYEEFVQMLTAKGGGGS
dNIRGECO0.8   FRVFDKDGNGYIGAAELRHVMTDLGEKLTDEKVDEMIRVADNDGDGQVNYEEFVQMLTAKGGGGS
dNIRGECO1.0   FRVFDKDGNGYIGAAELRHVMTDLGEKLTDVKVDEMIRVADNDGDGLVNYEEFVQMLTAKGGGGS

NIRGECO2G     VDSSRRKWNKAGHAVRAIGRLGSRRHDLLSECRRADLEAFLGNRYPASTIPQIARRLYELNRVRL  390
dNIRGECO0.2   VDSSRRKWNKAGHAVRTIGRLGSRRHDLLSECRRADLEAFLGNRYPASTIPQIARRLYELNRVRL
dNIRGECO0.4   VDSSRRKWNKAGHAVRTIGRLGSEPHDLLSECRRADLEAFLGNRYPASTIPQIARRLYELNRVRL
dNIRGECO0.6   VDSSRRKWNKAGHAVRTIGRLGSEPHGLLSECRRADLEAFLGSRYPASTIPQNARRLYELNRVRL
dNIRGECO0.8   VGSSRRKWNKAGHAVRTIGRLGSEPHGLLSECRRADLEAFLGSRYPASTIPQNARRLYELNRVRL
dNIRGECO1.0   VGSSRRKWNKAGHAVRTIGRLGSEPHGLLSECRRADLEAFRGSSYPASTIPNNARRLYELNRVRL

NIRGECO2G     LVDVNYTPVPLEPRISPLNGRDLDMSLSCLRSMSPIHQKYMQDMGVGATLVCSLMVSGRLWGLIV  455
dNIRGECO0.2   LVDVNCTPVPLEPRISPLNGRDLDMSLSCLRSMSPIHQKYMQDMGVGATLVCSLMVSGRLWGLIA
dNIRGECO0.4   LVDVNCTPVPLEPRISPLNGRDLDMSLSCLRSMSPIHQKYMQDMGVGATLVCSLMVSGRLWGLIA
dNIRGECO0.6   LVDVNCTPVPLEPRISPLNGRDLDMSLSCLRSMSPIHQKYMQDMGVGATLVCSLMVSGRLWGLIA
dNIRGECO0.8   LVDVNCTPVPLEPRISPLNGRDLDMSLSCLRSMSPIHQKYMQDMGVGATLVCSLMVSGRLWGLIA
dNIRGECO1.0   LVDVNCTPVPLEPRISPLNGRDLDMSLSCLRSMSPIHQKYMQDMGVGATLVCSLMVSGRLWGLIA

NIRGECO2G     CHHYEPRYVPSHIRAAGEALAEACANRIATLESFAQSQSK  480
dNIRGECO0.2   CHHYEPRYVPFHIRAAGEALAEACANRIATLESFAQSQSK
dNIRGECO0.4   CHHYEPRYVPFHIRAAGEALAEACANRIATLESFAQSQSK
dNIRGECO0.6   CHHYEPRYVPFHIRAAGAALAEACANRIATLESFAQSQSK
dNIRGECO0.8   CHHYEPRYVPFHIRAAGAALAEACANRIATLESFAQSQSK
dNIRGECO1.0   CHHYEPRYVPFHIRAAGAALAEACANRIATLESFAQSQSK
```

**Supplementary Figure 1.** Sequence alignment of NIR-GECO2G, dNIR-GECO0.2, dNIR-GECO0.4, dNIR-GECO0.6, dNIR-GECO0.8, and dNIR-GECO1.0. The calmodulin (CaM) domain is highlighted in cyan, RS20 peptide in green, mIFP in magenta, and linker regions in grey and black. Mutations relative to NIR-GECO2G are indicated in yellow.

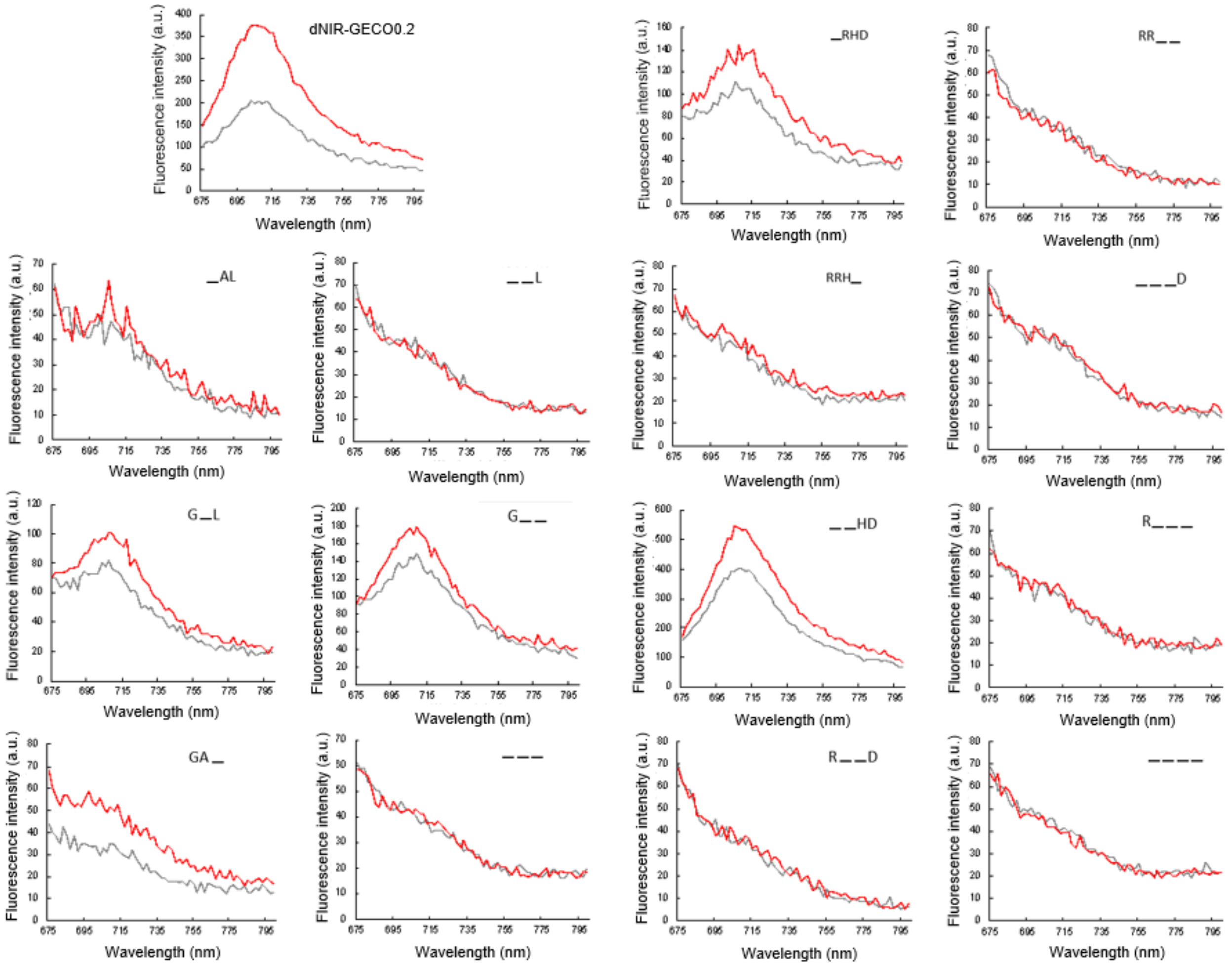

**Supplementary Figure 2.** Fluorescence emission spectra of the linker-deletion variants generated from dNIR-GECO0.2.

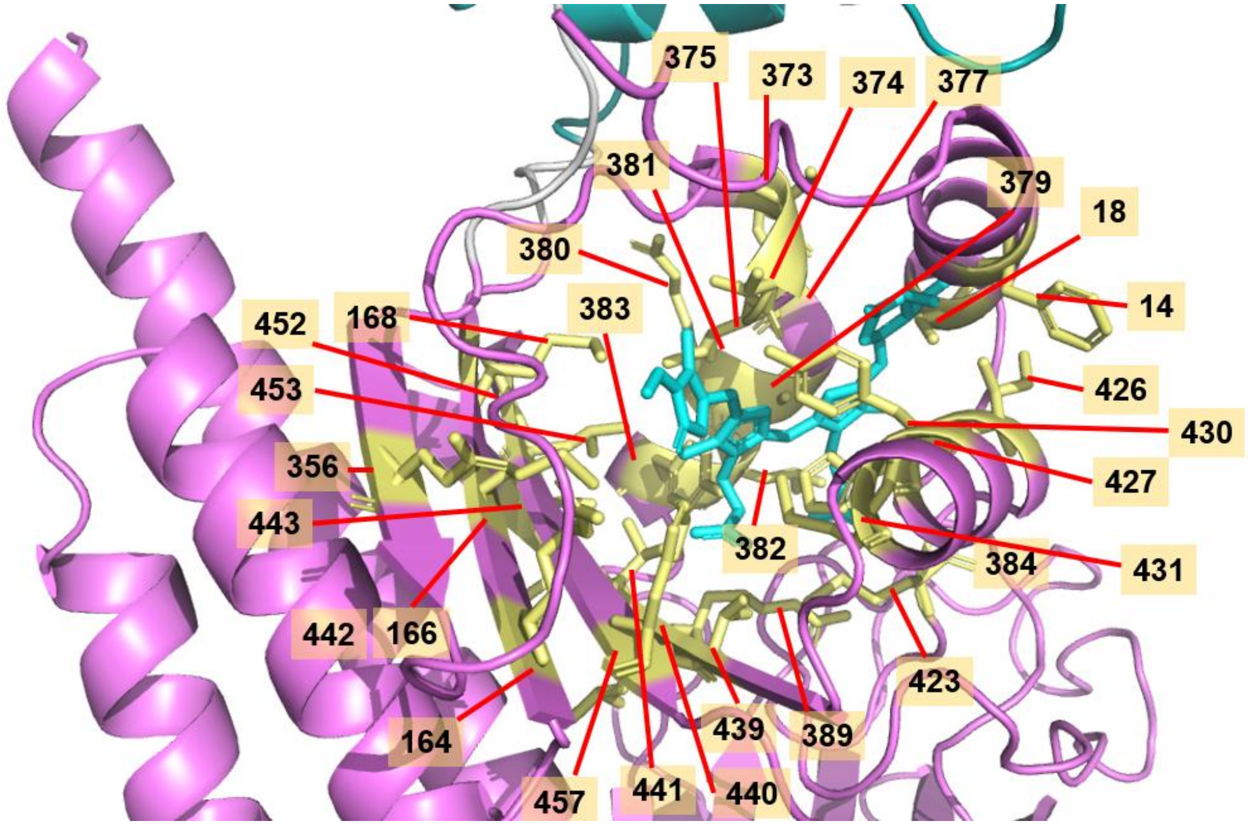

**Supplementary Figure 3.** Residues surrounding BV-binding pocket targeted for site-saturation mutagenesis during directed evolution.

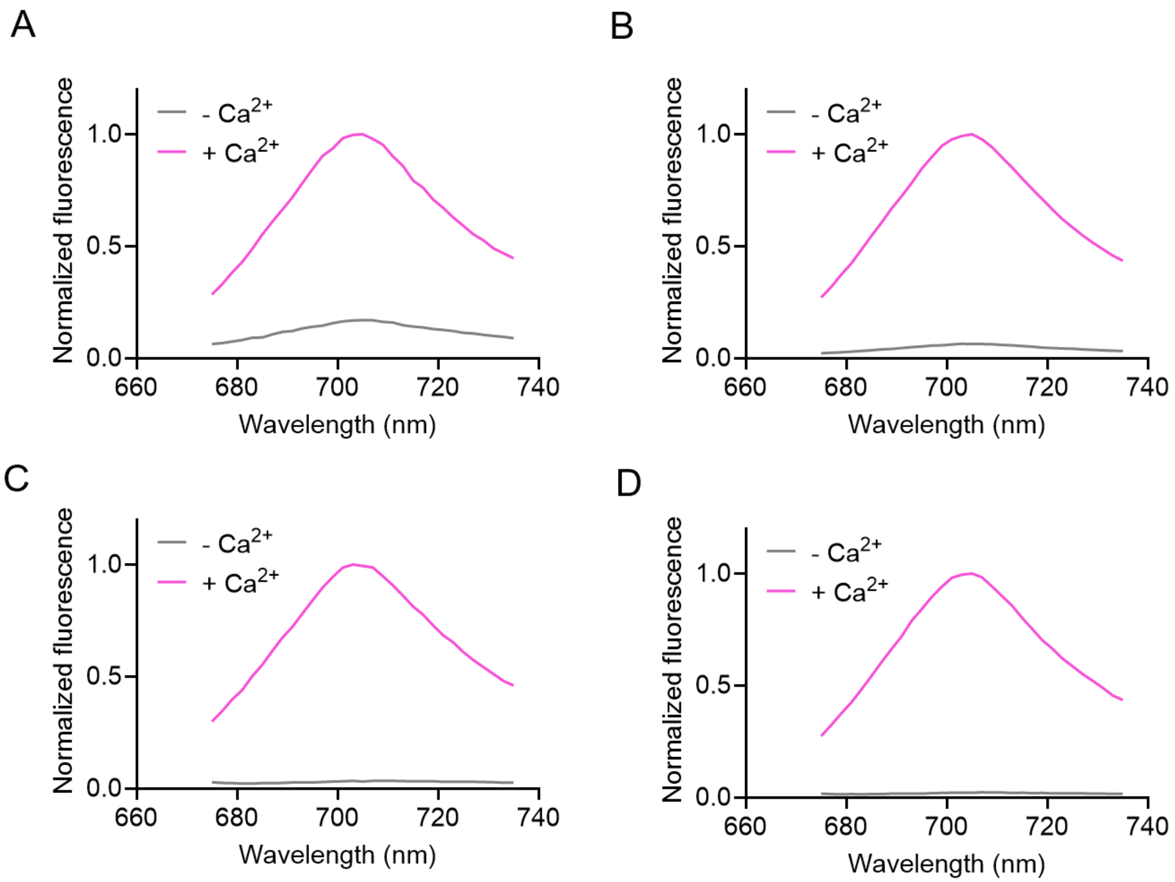

**Supplementary Figure 4.** Representative pH-dependent emission spectra of dNIR-GECO1.0 under different pH conditions. Emission spectra of dNIR-GECO1.0 recorded across acidic to basic conditions. (A) pH 5, (B) pH 7, (C) pH 9 and (D) pH 10. Spectra were normalized to the maximum emission intensity to compare pH-dependent spectral changes.

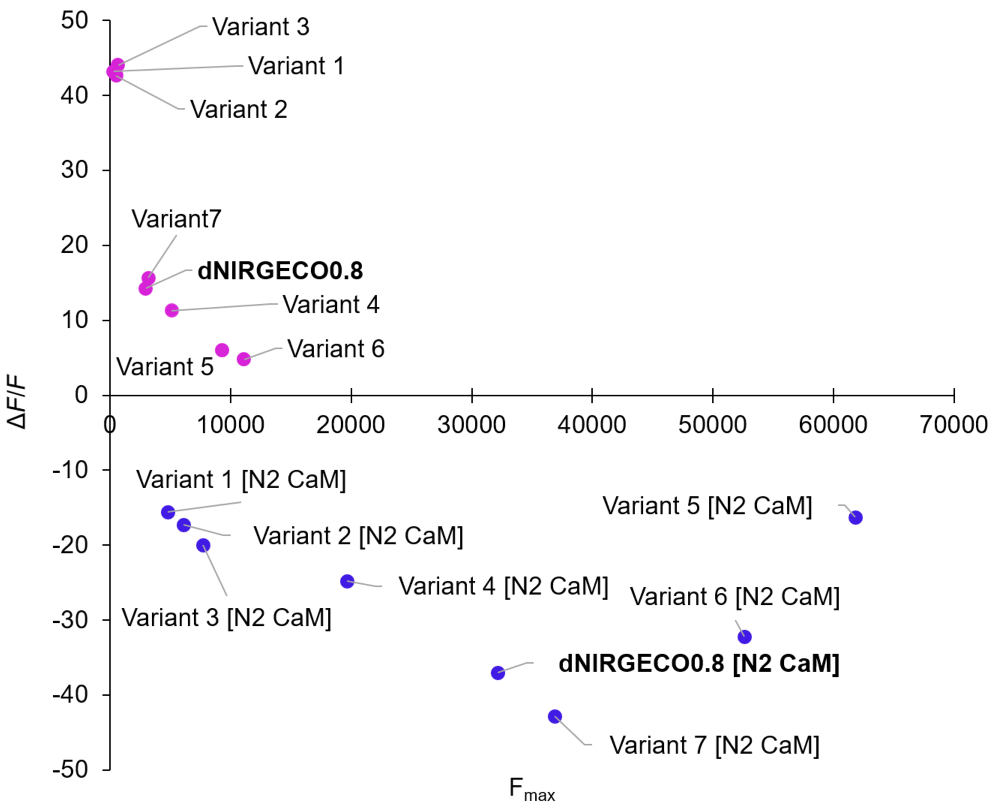

**Supplementary Figure 5.** Δ*F*/*F* versus $F_{max}$ of selected dNIR-GECO0.8 variants and corresponding variants containing the parental NIR-GECO2G CaM domain (N2 CaM). Pink data points indicate variants with the evolved CaM domain, whereas blue data points indicate variants with the parental CaM domain.

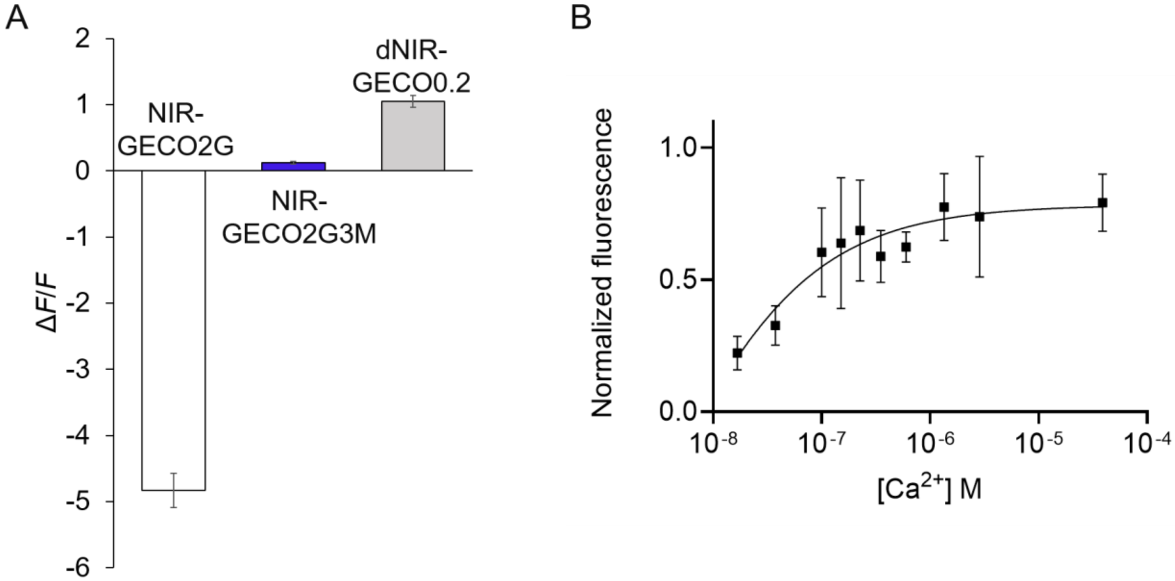

**Supplementary Figure 6.** Effect of three key mutations on direct response. (A) Fluorescence response upon $Ca^{2+}$ addition of NIR-GECO2G, NIR-GECO2G3M and dNIR-GECO0.2. (B) $Ca^{2+}$-dependent fluorescence titration of NIR-GECO2G3M (mean ± SD, n = 3).

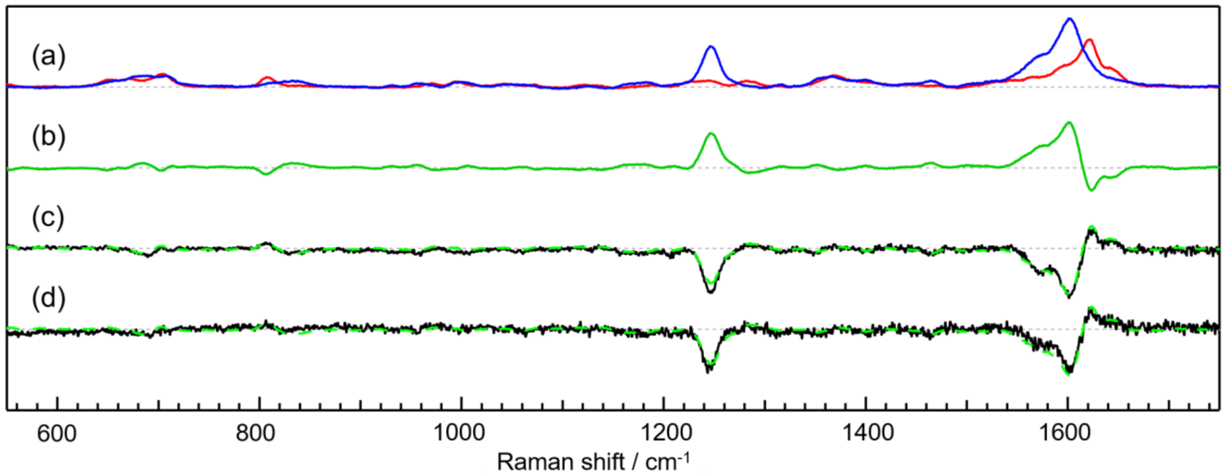

**Supplementary Figure 7.** Changes in the RR spectra of NIR-GECOs upon $Ca^{2+}$ binding. (a) RR spectra of NIR-GECO2G. The red and blue traces display the spectra in the bright and dimmer fluorescent state, respectively. The fluorescence background has been subtracted from each spectrum. (b)–(d) Difference spectra of (b) NIR-GECO2G (green), (c) dNIR-GECO1.0 (black), and (d) dNIR-GECO3M (black) upon $Ca^{2+}$ binding to the proteins. The green broken lines represent mirror relationships of the difference spectrum of NIR-GRCO2G, which were normalized by the intensity of the 1600 $cm^{-1}$ negative peak. The spectral contributions of impurities that could not be removed during purification have been canceled out.

### Supplementary Tables

**Supplementary Table 1.** Photophysical properties of dNIR-GECO variants.

| Protein | $Ca^{2+}$ | $\lambda_{max}$ abs (nm) | $\lambda_{max}$ em (nm) | ε ($mM^{-1}$ $cm^{-1}$) | QY | Molecular brightness ($mM^{-1}cm^{-1}$) | Molecular brightness fold increase (ε*QY) | p*K*a | Δ*F*/*F* Purified protein | $K_d$ (nM) | Hill coefficient |
|---|---|---|---|---|---|---|---|---|---|---|---|
| NIRGECO2G[a] | -ve | 680 | 703 | 74 | 0.06 | 4.51 | 10.3 | 5.34 | 6.6 | 480 | 0.78 |
| | +ve | 685 | 703 | 21 | 0.02 | 0.44 | | 4.84 | | | |
| GAF-CaMP2-sfGFP[b] | -ve | 630 | 676 | 15.7 | 0.03 | 0.52 | 3.7 | 5.3, 7.3 | 0.9 | 289 | NA |
| | +ve | 649 | 674 | 27.5 | 0.07 | 1.90 | | 4.9, 7.6 | | | |
| GAF-CaMP3-sfGFP[b] | -ve | 640 | 674 | 14.4 | 0.05 | 0.72 | 2.9 | 3.5, 9.2 | 2.0 | 433 | NA |
| | +ve | 654 | 676 | 27 | 0.08 | 2.08 | | 3.5, 8.9 | | | |
| dNIR-GECO0.2 | -ve | 685 | 704 | 21.3 | 0.04 | 0.85 | 1.6 | 5.03, 7.74 | 1.1 | 101 | 1.7 |
| | +ve | 680 | 704 | 27.1 | 0.05 | 1.35 | | 6.14, 6.19 | | | |
| dNIR-GECO0.6 | -ve | 685 | 704 | ND | ND | ND | ND | ND | 8.0 | 112 | 2.0 |
| | +ve | 678 | 704 | ND | ND | ND | | | | | |
| dNIR-GECO0.8 | -ve | 685 | 704 | 12.8 | 0.03 | 0.38 | 7.6 | 5.00, 6.05 | 12.1 | 103 | 2.0 |
| | +ve | 678 | 704 | 35.8 | 0.08 | 2.87 | | 5.06, 9.06 | | | |
| dNIR-GECO1.0 | -ve | 685 | 704 | 17.1 | 0.02 | 0.34 | 18.1 | 5.00, 6.02 | 22.1 | 181 | 1.4 |
| | +ve | 679 | 704 | 68.4 | 0.09 | 6.16 | | 5.05, 9.67 | | | |

Abbreviations: -ve, negative; +ve, positive; $\lambda_{max}$ abs, absorbance maximum; $\lambda_{max}$ em, fluorescence emission maximum; ε, extinction coefficient; QY, quantum yield; p$K_a$, pH corresponding to 50% of the maximal fluorescence brightness measured at optimal pH; $K_d$, dissociation constant for $Ca^{2+}$, ND, not determined; NA, not available.

[a]Values from Qian *et al.*[15]

[b]Values from Subach *et al.*[12]

**Supplementary Table 2.** Sequence of linker regions and fluorescence property of top variants from each round of linker optimization.

| **Round** | **Variant** | **1st linker region** | **2nd linker region** | **Δ*F*/*F*** | **Fluorescence intensity** |
|---|---|---|---|---|---|
| | dNIR-GECO0.2 | MVMRL**GAL**DDVTE | GRLGS**RRHD**LLSEC | 1.63 | 1 |
| 1st | dNIR-GECO0.2-1 (**dNIR-GECO0.2**) | MVMRL**GAL**DDVTE | GRLGS**RRHD**LLSEC | 1.63 | 1 |
| 2nd | dNIR-GECO0.2-2 | MVMRL**GKL**DDVTE | GRLGS**RRHD**LLSEC | 1.93 | +14% than dNIR-GECO0.2-1 |
| 3rd | dNIR-GECO0.2-3 | MVMRL**GAL**DDVTE | GRLGS**RPHD**LLSEC | 1.61 | +156% than dNIR-GECO0.2-2 |
| 4th | dNIR-GECO0.2-4 (**dNIR-GECO0.4**) | MVMRL**GAL**DDVTE | GRLGS**EPHD**LLSEC | 1.35 | +80% than dNIR-GECO0.2-3 |

**Supplementary Table 3.** *In vitro* photophysical characterization of selected high-response variants identified during directed evolution.

| Protein | [$Ca^{2+}$] (μM) | λmax abs (nm) | λmax em (nm) | ε ‘ ($mM^{-1}cm^{-1}$) | Φ | Molecular brightness ($mM^{-1}cm^{-1}$) | Molecular brightness fold increase (ε*QY) | Δ*F*/*F* |
|---|---|---|---|---|---|---|---|---|
| HRV1 | 0 | 685 | 703 | 14.27 | 0.01 | 0.14 | 23.1 | 38.2 |
| | 39 | 678 | 703 | 49.75 | 0.07 | 3.48 | | |
| HRV2 | 0 | 680 | 703 | 13.15 | 0.01 | 0.13 | 28.7 | 58.8 |
| | 39 | 685 | 703 | 48.77 | 0.08 | 3.9 | | |
| HRV3 | 0 | 685 | 703 | 14.34 | 0.01 | 0.14 | 27.3 | 44.3 |
| | 39 | 678 | 703 | 50.6 | 0.08 | 4.05 | | |

**Supplementary Table 4.** Mutational analysis of mIFP and CaM domains in NIRGECO2G. Columns are colored according to domain: magenta, mIFP; cyan, CaM; and green, RS20.

| **Variants** | | **NIR-GECO2G** | **NIRGECO2G +** | | | | | | | | | | | **dNIR-GECO0.2** |
|---|---|---|---|---|---|---|---|---|---|---|---|---|---|---|
| | | | **I77V** | **K124E** | **L176V** | **G197R** | **A218V** | **D228Y** | **E292K** | **A342T** | **Y396C** | **V455A** | **S466F** | |
| ***ΔF/F*** | 120 nM $Ca^{2+}$ | -0.3 | -0.3 | -0.3 | -0.1 | 0.0 | -0.3 | -0.1 | -0.2 | 0.0 | -1.1 | -0.5 | -0.3 | 0.6 |
| | 39 μM $Ca^{2+}$ | -7.3 | -6.6 | -7.7 | -4.3 | -2.9 | -7.0 | -0.4 | -6.6 | -1.7 | -45.6 | -30.3 | -11.0 | 0.7 |
| | 0.5 mM $Ca^{2+}$ | -7.7 | -7.3 | -8.4 | -4.9 | -5.8 | -7.6 | -1.1 | -7.2 | -2.3 | -47.4 | -31.2 | -12.1 | 0.6 |

**Supplementary Table 5.** List of variants selected for optimization of the CaM domain. Columns are colored according to domain: magenta, mIFP; cyan, CaM; and green, RS20.

| **dNIR-GECO0.8 +** | **F10Y** | **A182E** | **E291V** | **Q307L** | **S329T** | **L366R** | **R369S** | **Q377N** | **M423L** | **T447S** | **V455A** | **C463Y** | **A473E** |
|---|---|---|---|---|---|---|---|---|---|---|---|---|---|
| Variant 1 (HRV1) | | / | | | | | | | | / | / | / | / |
| Variant 2 (HRV2) | | / | | | / | | | | / | / | / | / | / |
| Variant 3 (HRV3) | / | / | | | | | | | / | / | / | / | / |
| Variant 4 | | / | | | | | / | | | | | | |
| Variant 5 | | | | | | | / | / | | | | | |
| Variant 6 | | | | | | | | / | | | | | |
| Variant 7 | | | / | / | | / | / | / | | | | | |

**Supplementary Table 6.** Mutational analysis of the CaM domain in dNIR-GECO0.8 using the parental NIR-GECO2G CaM (N2 CaM). The evolved CaM domain of dNIR-GECO0.8 was replaced with the parental NIRGECO2G CaM (N2 CaM) and individual mutations were reintroduced one at a time and characterized for $Ca^{2+}$-dependent analysis. Columns are colored according to domain: cyan, CaM; and green, RS20.

| Variants | | dNIR-GECO0.8 | dNIR-GECO0.8 [N2 CaM] + | | | | | | | | | | | |
|---|---|---|---|---|---|---|---|---|---|---|---|---|---|
| | | | 176 | 197 | 208 | 218 | 228 | 238 | 243 | 246 | 292 | 317 | 327 | 342 |
| $F_{max}$ (0 mM $Ca^{2+}$) | | 30628.3 | 34907 | 30182 | 25407 | 4339 | 49893 | 35945 | 22092 | 23597 | 2490 | 17878 | 21210 | 18468 |
| $F_{max}$ (120 nM $Ca^{2+}$) | | 49401 | 55964 | 49443 | 41923 | 8733.7 | 55138 | 54168 | 51733 | 43605 | 20976 | 35238 | 33394 | 44401 |
| Δ*F*/*F* | 120 nM $Ca^{2+}$ | 0.6 | 0.6 | 0.6 | 0.7 | 1 | 0.08 | 0.5 | 1.3 | 0.8 | 7.4 | 1 | 0.6 | 1.4 |
| | 39 μM $Ca^{2+}$ | -46.8 | -22.2 | -3.1 | -36 | -22.6 | 0.03 | -39.6 | -1.2 | -25.8 | -3.7 | -38.9 | -47.2 | -5 |
| | 0.5 mM $Ca^{2+}$ | -62.5 | -43.6 | -29 | -59.9 | -29.3 | -0.8 | -65.7 | -16.6 | -43.9 | -5.7 | -55.2 | -66.8 | -21 |

**Supplementary Table 7.** Mutational analysis of the CaM domain in dNIR-GECO0.8. Each evolved CaM mutation in dNIR-GECO0.8 was individually reverted, and the resulting variants were characterized for $Ca^{2+}$-dependent fluorescence responses. Columns are colored according to domain: cyan, CaM; and green, RS20.

| Variants | | dNIR-GECO0.8 | dNIR-GECO0.8 - | | | | | | | | | | | |
|---|---|---|---|---|---|---|---|---|---|---|---|---|---|---|
| | | | **176** | **197** | **208** | **218** | **228** | **238** | **243** | **246** | **292** | **317** | **327** | **342** |
| **$F_{max}$ (39 µM $Ca^{2+}$)** | | 36849 | 17686 | 15735 | 41551 | 49950 | 43614 | 14544 | 39951 | 43184 | 59696 | 42228 | 44287 | 44287 |
| ***ΔF/F*** | **120 nM $Ca^{2+}$** | 5.8 | 7.4 | 5.8 | 4.8 | 5.8 | 5.5 | 6.6 | 5.8 | 5.3 | 4.3 | 7.1 | 6.3 | 7.5 |
| | **39 µM $Ca^{2+}$** | 12.9 | 17.6 | 10.6 | 15.1 | 13.6 | 12.6 | 16.1 | 13.7 | 15.2 | 5.1 | 12.4 | 12.2 | 10.7 |
| | **0.5 mM $Ca^{2+}$** | 12.8 | 16.9 | 9.8 | 14.9 | 14.1 | 10.1 | 16.3 | 13.7 | 14.8 | 4.8 | 12.0 | 11.8 | 10.6 |

**Supplementary Table 8.** Pairwise combination analysis of CaM mutations in dNIR-GECO0.8. Columns are colored according to domain: cyan, CaM; and green, RS20.

| **Variants** | **Mutations** | | | | | | | | **[$Ca^{2+}$]** | | |
|---|---|---|---|---|---|---|---|---|---|---|---|
| | **176** | **197** | **228** | **243** | **246** | **292** | **327** | **342** | **120 nM** | **39 µM** | **0.5 mM** |
| dNIR-GECO0.8 | | | | | | | | | 6.6 | 14.9 | 14.5 |
| dNIR-GECO0.2 | | | | | | | | | 0.7 | 0.9 | 0.7 |
| dNIR-GECO0.8 [N2 CaM] | | | | | | | | | 0.6 | -43 | -62.4 |
| dNIR-GECO0.8 [N2 CaM] + | | | | | / | | | | 0.8 | -25.8 | -43.9 |
| | | | | | | / | | | 7.4 | -3.7 | -5.7 |
| | | | / | | | | | | 0.1 | 0.04 | -0.5 |
| | | | | | | | / | | 0.6 | -47.2 | -66.8 |
| | | | | | | | | / | 1.4 | -5 | -21 |
| | / | | | | | | | | 0.6 | -22.2 | -43.6 |
| | | | | / | | | | | 1.3 | -1.2 | -16.6 |
| | | / | | | | | | | 0.6 | -3.1 | -29 |
| | | | / | | | / | | | 5.6 | 5.6 | 1.7 |
| | | | | / | | / | | | 8 | 3.5 | -1 |

| | | | | | | | | | | | |
|---|---|---|---|---|---|---|---|---|---|---|---|
| | | | | | | / | / | | 6 | -3.9 | -8.2 |
| | | | | | | | / | / | 1.6 | -2.5 | -19.6 |
| | | | / | / | | | | | 0.7 | 0.6 | 0.1 |
| | / | | | / | | | | | 0.7 | -0.2 | -9.2 |
| | | | | | | / | | / | 8.5 | 0.6 | -2.2 |
| | | / | / | | | | | | 0.7 | 0.8 | 0.7 |
| | | | / | | | | | / | 1.3 | 1.2 | -0.2 |
| | | | | / | | | | / | 2.4 | 0 | -11.8 |
| | | / | | | | / | | | 6 | 2.3 | 2.2 |
| | | / | / | / | | | | | 0.9 | 1 | 0.9 |
| | | / | / | | | / | | | 6.4 | 7.9 | 7.9 |
| | | / | / | | | | | / | 1.6 | 1.8 | 1.7 |
| | | | / | / | | | | / | 1.5 | 1.4 | -0.2 |
| | | | / | | | / | | / | 8.2 | 11.8 | 5.9 |
